# SLC11A2 affects nutritional immunity in the gut epithelium

**DOI:** 10.1101/2025.11.12.688034

**Authors:** Emilia S. Norberg, Trina L. Westerman, Eddy Cruz, Summer D. Bushman, Eric P. Skaar, Johanna R. Elfenbein, Leigh A. Knodler

**Author notes:** Corresponding authors: Leigh A. Knodler, and Johanna R. Elfenbein.

## Abstract

There is a constant tug-of-war for transition metals at the pathogen-host interface. A goal of the vertebrate host is to modulate the availability of metals to pathogens, in a process known as nutritional immunity, but pathogens have evolved numerous countermeasures to regulate intracellular trace metal levels. The bioavailability of trace metals therefore shapes the outcome of disease. In the human body, epithelial cells lining the intestine are a major site of metal absorption. Intestinal epithelial cells (IECs) are also a target for invading enteric pathogens but the contribution of epithelium-intrinsic factors towards nutritional immunity has been understudied. Using *Salmonella enterica* serovar Typhimurium (STm) harboring metal-responsive fluorescent reporters in a bovine ligated intestinal loop infection model, we mapped the spatiotemporal nature of metal competition during enteric salmonellosis. We show that STm experiences a temporal, cell-specific restriction of iron, manganese, and zinc in the intestinal mucosa during the early stages of infection. We have further studied the contribution of the broad specificity metal cation transporter, SLC11A2, in IECs to nutritional immunity against STm. Knockout of SLC11A2 in IECs leads to enhanced replication of STm, indicating a protective role for this transporter. Using fluorescence-based biosensors and bacterial gene deletion mutants, we pinpoint manganese and iron restriction as the mechanism by which SLC11A2 limits bacterial proliferation. We conclude that SLC11A2-mediated sequestration of metals is an intrinsic defense mechanism of the intestinal epithelium against enteric bacteria.

**Significance statement:** There is limited source of trace metals in the gut that invading pathogens and the infected host must compete for. Using *Salmonella enterica* as a model enteric pathogen, along with fluorescent reporters that respond to metal ion availability, we traced the sites of metal ion limitation in the intestinal mucosa to intestinal epithelial cells (IECs) and phagocytes in the underlying lamina propria. We further show that SLC11A2 (NRAMP2), which is the sole SLC11 family member expressed in IECs and localizes to the apical membrane and endosomal network, limits the intracellular proliferation of *Salmonella enterica* by withholding iron and manganese. Therefore SLC11A2-mediated nutritional immunity is an IEC-intrinsic defense mechanism that protects against microbial pathogens.

## Introduction

Pathogens and their vertebrate hosts both require trace nutrient metals for cellular functions. To establish infection, pathogenic microorganisms actively scavenge trace nutrient metals from the extracellular space and host cells plus possess mechanisms to minimize metal toxicity. As a counter-defense, the host can withhold transition metals to starve, or deliver metals in excess to intoxicate, microbial invaders. This concept is known as “nutritional immunity” (1, 2). The solute carrier 11 (SLC11) or <u>N</u>atural resistance-<u>a</u>ssociated <u>m</u>acrophage <u>p</u>rotein (NRAMP) family sits at the nexus between host and microbe in the tug-of-war for metals. These transition metal transporters are conserved from bacteria to humans. In bacteria, SLC11/NRAMP proteins are represented by the proton-dependent manganese transporter (MntH) family. There are two SLC11 paralogs - *SLC11A1* (also known as *NRAMP1*) and *SLC11A2* (also known as *NRAMP2*, <u>d</u>ivalent <u>m</u>etal transporter 1 (*DMT1*) or <u>d</u>ivalent <u>c</u>ation transporter 1 (*DCT1*)) - in humans, and *Slc11a1* and *Slc11a2* in mice. Both are multi-spanning membrane proteins. While SLC11A1 is a pH-dependent antiporter of Fe^2+^, Mn^2+^ and Zn^2+^ (3), SLC11A2 is a H^+^/metal ion symporter with broader substrate specificity (3–7). SLC11A2 also transfers iron from endosomes into the cytosol after transferrin-bound iron is endocytosed by CD71, alternatively known as transferrin receptor 1 (TfR) (8).

*Slc11a1* was originally identified as a locus that confers natural resistance of some inbred mice strains to *Mycobacterium, Leishmania* and *Salmonella* infection (9–11). These inbred mice have a naturally occurring missense mutation (G169D) in Slc11a1 (12, 13) and are characterized by the absence of the mature glycosylated protein form of the transporter (14, 15). *Slc11a1*^G169D^ and *Slc11a1*^-/-^ mice are phenotypically indistinguishable (12). Natural resistance to STm has also been linked with sequence variations in *Slc11a1* in chickens (16). In humans, allelic variants of *SLC11A1* are associated with susceptibility to tuberculosis, leprosy and leishmaniasis (17–19). *Slc11a1* expression is restricted to myeloid cells, *e.g.* macrophages, dendritic cells and neutrophils (20–22), and control of intracellular microbial growth is thought to be via depletion of transition metals from phagolysosomes. However, it remains controversial as to which divalent metals are restricted within the phagosome (23–26). Slc11a1/SLC11A1 also increases pro-inflammatory anti-microbial immune responses via nitric oxide and lipocalin production, bactericidal activity of neutrophils, and activation of innate lymphocytes (27–30).

Unlike *Slc11a1*, *Slc11a2*/*SLC11A2* is ubiquitously expressed and particularly abundant in the proximal intestine ((4); The Human Protein Atlas; https://www.proteinatlas.org/ENSG00000110911-SLC11A2/tissue). Slc11a2 was first identified as a divalent metal ion transporter in inbred mice (4, 31) and rats (8). Microcytic anemia (*mk/mk*) mice and Belgrade (*b/b*) rats are iron and manganese deficient (32–34) due to a point mutation (G185R) in Slc11a2 which affects its stability and trafficking (35), and reduces its transport of Fe^2+^, Ni^2+^ and Mn^2+^ (32, 36, 37). The *mk/mk* mouse and *b/b* rat anemia phenotypes are, however, less severe than that of *Slc11a2*^-/-^ mice, which die soon after birth (38). SLC11A2 mutations are extremely rare in humans; the few reported impact exon splicing (39, 40), or lead to point mutations (41–43) or amino acid deletions (41) in SLC11A2. Patients exhibit microcytic anemia, and hepatic iron overload appears at a young age in some cases (39, 40, 42, 43). *SLC11A2* expression and/or SLC11A2 function have also been implicated in the pathogenesis of neurodegenerative diseases and inflammatory bowel disorders (44–46). The impact of SLC11A2 on infection outcome is not known.

*Salmonella enterica* infection in humans presents as distinct clinical syndromes depending upon the serotype. Non-typhoidal *Salmonella* (NTS) serotypes, e.g. *S*. Typhimurium (STm) and *S*. Enteritidis, cause self-limiting diarrhea in immunocompetent individuals characterized by severe neutrophilic intestinal inflammation, or an invasive disease characterized by bacteremia in immunocompromised individuals. During typhoid fever, a systemic disease caused by human host restricted *S*. Typhi and *S*. Paratyphi, bacteria persist within professional phagocytes in granulomas in the liver, spleen and bone marrow (47). Although the clinical syndromes differ between serotypes, the intestinal epithelium is the first line of host defense against these food-borne pathogens.

*In vivo* studies have convincingly defined the importance of Slc11a1 in myeloid cells to nutritional immunity and, as a result, Slc11a1 has assumed a place of prominence in metal deprivation during infection. However, *Slc11a1-*mediated restriction of bacterial growth is temporal, tissue-specific and depends on the infection route in murine models (22, 48). For example, despite the presence of Slc11a1-positive cells in the intestinal lamina propria, and their enrichment during STm infection (22, 49), *Slc11a1* has little to no effect on STm replication in the mouse gut (48, 49). By contrast, the primary sites of *Slc11a1*-dependent control against STm are the spleen, liver and mesenteric lymph nodes where STm primarily resides within macrophages (24, 48, 49). Altogether, these observations raise the question of whether other SLC11 proteins also contribute to host defense. To close this knowledge gap, here we sought to define the relationship between SLC11A2, intestinal epithelial cells (IECs), and nutritional immunity during infection with the model enteric pathogen, STm. Using a natural host for NTS, the calf enteritis model (50), in combination with STm harboring metal ion responsive reporters, we show that the mammalian host restricts iron, manganese and zinc bioavailability in the intestinal mucosa, first in IECs, then in lamina propria phagocytes. Further, using CRISPR/Cas9 edited IECs we establish that SLC11A2 deprives iron and manganese from intracellular STm to limit their proliferation. Therefore, SLC11A2-mediated nutritional immunity is an IEC-intrinsic defense mechanism against invading pathogens.

## Results

### STm is starved of trace metals during the acute stages of enteric infection

The calf is a relevant animal model for studying acute *Salmonella*-induced enteritis because STm is a natural pathogen of cattle, STm infection causes similar signs of disease and pathology as observed in humans (50) and the *SLC11A1* G169D mutation is not found in European breeds of cattle (51). The ligated ileal-jejunal loop model in calves is especially suited for investigating *Salmonella*-host interactions in the gut at early times post-infection (50). To validate this model for the study of SLC11-mediated metal ion restriction during acute enteritic infection, we immunolocalized SLC11A1 and SLC11A2 in uninfected intestinal tissue. SLC11A1 localized to discrete cells in the lamina propria and submucosa (Figure 1), likely phagocytes and endothelial cells given that bovine *SLC11A1* is primarily expressed in these cell types (29, 52). SLC11A1 is similarly restricted to lamina propria phagocytes in the human colon (The Human Protein Atlas; https://www.proteinatlas.org/ENSG00000018280-SLC11A1/tissue) and mouse small intestine (22, 49). By contrast, SLC11A2 staining was concentrated at the luminal surface of IECs as well as puncta within IECs in the calf ileum (Figure 1), like for human duodenum (53–55) and colon (The Human Protein Atlas; https://www.proteinatlas.org/ENSG00000110911-SLC11A2/tissue). The distinct anatomic and subcellular distributions of these two metal transporters in the mammalian gastrointestinal tract suggest that they play discrete roles in host defense.

**Figure 1:**
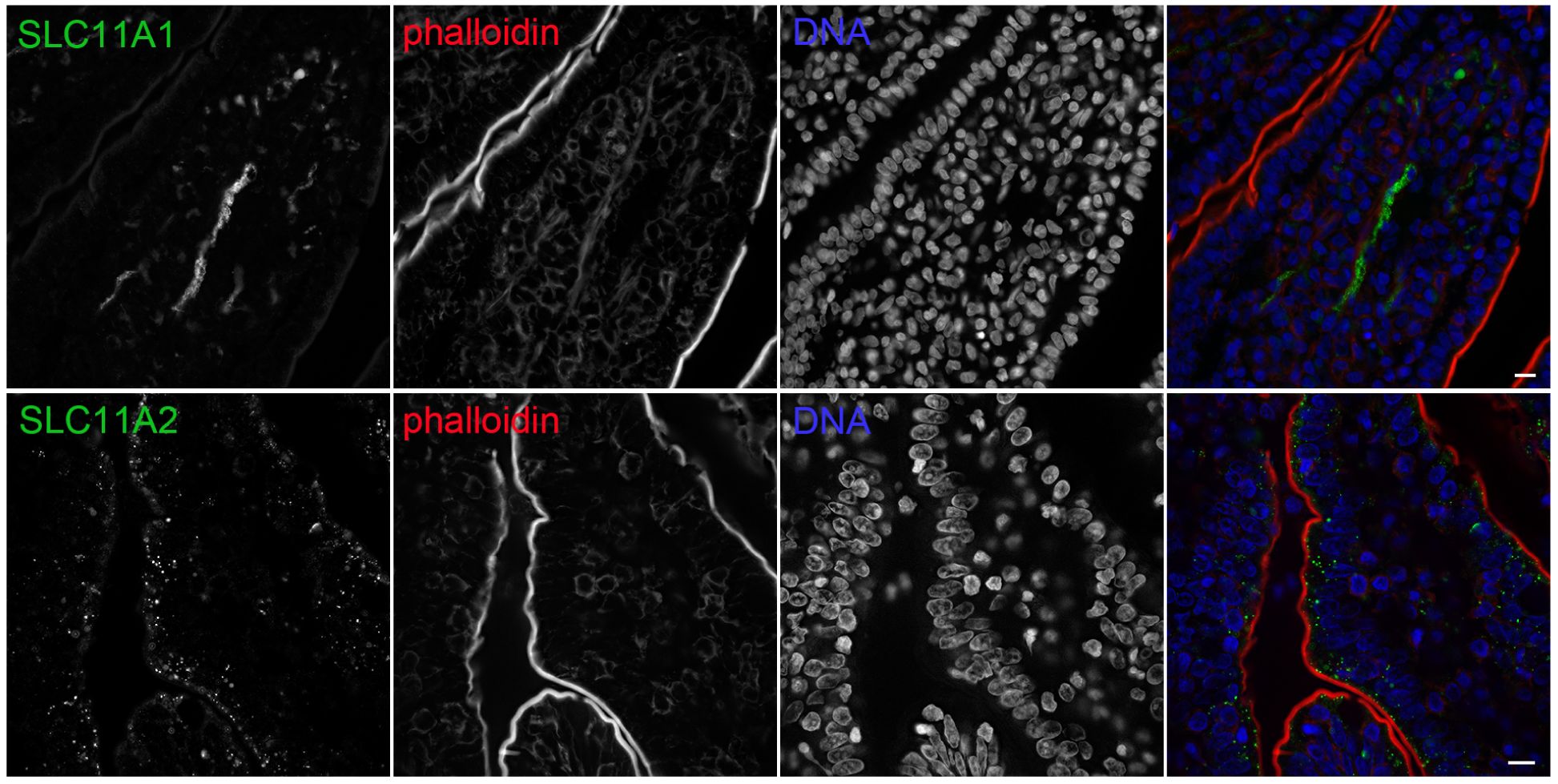
Spatially distinct localization of SLC11A1 and SLC11A2 in intestinal mucosa. Mock-infected (2 h post-inoculation with PBS) bovine intestinal tissues were immunostained with polyclonal antibodies against SLC11A1 (NRAMP1, upper panels) or SLC11A2 (NRAMP2, lower panels). Phalloidin was used to visualize F-actin at the apical surface of epithelial cells. DNA was stained with DAPI. Scale bars are 10 µm.

We next determined the spatiotemporal nature of metal ion availability during acute *Salmonella*-induced enteritis. We previously showed that STm carrying plasmids encoding *iroN*, *sitA* or *zinT* promoters fused to *gfpmut3* serve as sensitive reporters of iron, iron/manganese, and zinc concentrations, respectively, *in vitro* and *in cellulo* (56). Expression of these genes, *i.e.* GFP fluorescence, is induced when the respective metal ions are <1 µM and is maximal at <0.1 µM (56). *iroN* encodes a receptor for enterobactin and salmochelin, two high-affinity iron-binding siderophores (57–59), the *sitABCD* operon encodes an ABC-type manganese importer that belongs to the NRAMP superfamily (60, 61) and *zinT* encodes an accessory protein for the high-affinity zinc importer, ZnuABC (62). Bovine ileal loops were inoculated with wild-type bacteria constitutively expressing *mCherry* on the chromosome (STm-mCherry) and harboring P*iroN*-*gfpmut3*, P*sitA*-*gfpmut3* or P*zinT*-*gfpmut3* reporters. Tissues were collected at 2 h or 8 h post-inoculation, fixed, and sectioned. The infection times represent prior to (2 h), and after (8 h), severe neutrophilic inflammation (63). Tissue sections were stained with phalloidin to demarcate the epithelial brush border and DAPI for DNA (host cell nuclei). The number of tissue-associated GFP-positive STm-mCherry in the “epithelium” (*i.e.* bacteria in the single layer of cells lining the villi), or “lamina propria” (*i.e.* bacteria in the tissue beneath the epithelium), was scored by fluorescence microscopy (Figure 2A). GFP-positive bacteria were detected as early as 2 h post-inoculation – 17.0±7.0% for P*iroN*-*gfpmut3*, 38.7±19.5% for P*sitA*-*gfpmut3* and 14.0±6.7% for P*zinT*-*gfpmut3* – indicating that bacteria are deprived of iron, manganese and zinc shortly after internalization into the intestinal mucosa (Figure 2A). GFP-positive bacteria were primarily within IECs at 2 h (Figure 2A, 2B). By 8 h post-inoculation, 38.3±11.8% of P*iroN-gfpmut3*, 32.5±14.4% of P*sitA*-*gfpmut3*, and 7.6±6.1% of P*zinT*-*gfpmut3* harboring bacteria were GFP-positive (Figure 2A). Most of these were in the lamina propria (Figure 2A). At both 2 h and 8 h, more P*iroN*-*gfpmut3* and P*sitA*-*gfpmut3* bacteria were GFP-positive than P*zinT*-*gfpmut3* bacteria, hinting that bioavailability of iron and manganese is more limited than zinc in infected tissues. From these data we conclude that STm is starved for trace metals in multiple cell types and at distinct anatomic sites during the early stages of enteric infection.

**Figure 2:**
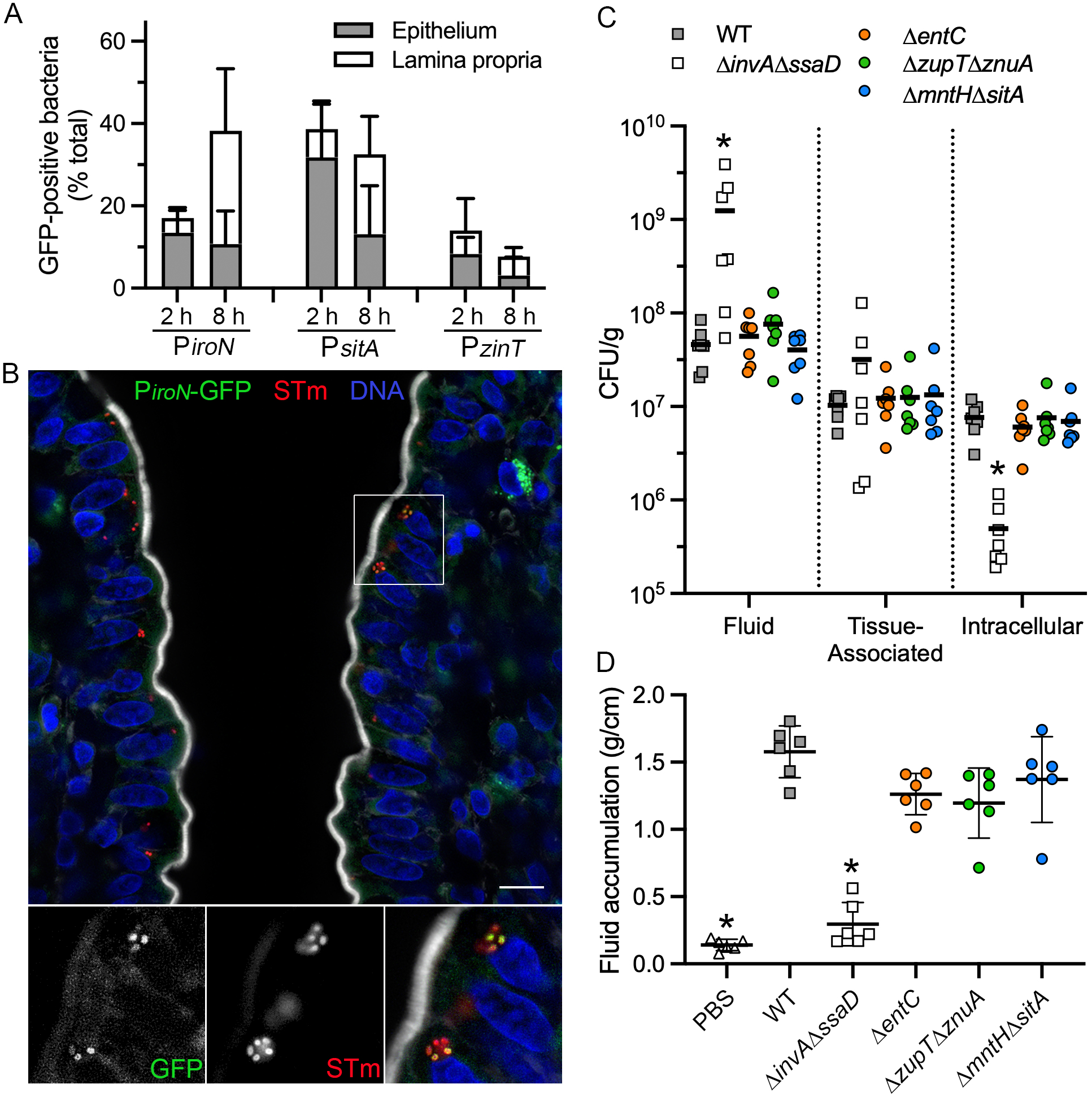
*Salmonella* are deprived of iron, manganese and zinc in the gut. (A) Ligated jejunal-ileal loops were inoculated with STm-mCherry harboring P*iroN*-*gfpmut3*, P*sitA*-*gfpmut3* or P*zinT*-*gfpmut3* transcriptional reporters (∼10^9^ CFU). GFP fluorescence intensity increases as levels of iron, iron/manganese, and zinc, respectively, become more limiting. Tissues were collected at 2 h or 8 h post-inoculation and fixed. Sections were stained with phalloidin to demarcate the apical surface of epithelial cells and Hoechst 33342 for DNA. Tissue-associated GFP-positive bacteria were scored within the epithelium (*i.e.* the single layer of cells lining the villi) or the lamina propria (*i.e.* in the tissue beneath the epithelium). Mean ± SD from n ≥ 4 calves. (B) Representative fluorescence microscopy image of intestinal tissue infected with STm-mCherry harboring P*iroN*-*gfpmut3* reporter at 2 h post-inoculation. GFP-positive bacteria are shown in green, all *Salmonella* are red, DNA is blue and actin is greyscale. Scale bar is 10 µm. (C and D) Ligated ileal loops were inoculated with PBS, wild-type (WT) bacteria or the indicated deletion mutant (∼10^9^ CFU) for 8 h. CFU from washed intestinal tissue (total tissue-associated bacteria), gentamicin-treated intestinal tissue (intracellular) and fluid were enumerated by serial dilution and plating on LB agar. CFU were normalized to the tissue or fluid weight (C). Fluid weight was normalized to loop length (D). Each symbol represents data from a single loop from one calf. *p<0.05, significantly different from ST4/74 WT bacteria, ANOVA with multiple comparisons.

We tested the impact of metal limitation on bacterial survival in the calf intestine. We compared the fitness of wild-type bacteria with Δ*entC* (defective in iron-binding siderophore synthesis, *i.e.*, enterobactin and salmochelin), Δ*mntH*Δ*sitA* (defective in Mn^2+^ >> Fe^2+^ import), and Δ*zupT*Δ*znuA* (defective in Zn^2+^ >> Mn^2+^ and Co^2+^ import) mutants at 2 h and 8 h post-inoculation. We observed no significant differences in bacterial burden for any metal mutant in any tissue compartment (Figures 2C and S1). There was a small but not statistically significant reduction in fluid accumulation, a proxy for neutrophilic inflammation (63), in loops inoculated with metal acquisition mutants at 8 h (Figure 2D). The non-invasive Δ*invA*Δ*ssaD* mutant, which lacks structural components of both the T3SS-1 (defective for tissue invasion and induction of neutrophilic inflammation) and T3SS-2 (defective for intracellular replication), induced very little fluid accumulation (Figure 2D) and was severely attenuated for intracellular colonization (Figure 2C), as expected (64–66). These data indicate that STm mutants deficient in manganese, iron and zinc import, and siderophore biosynthesis, are not defective for intestinal colonization during the first 8 h of enteric disease in calves.

### *SLC11A2* deletion leads to enhanced bacterial proliferation in IECs

Being as SLC11A2 is highly expressed in IECs (Figure 1), transports multiple metal ions into mammalian cells (33, 37, 67–69), and STm experiences transition metal limitation in IECs *in vivo* (Figure 2), we hypothesized that IEC-intrinsic SLC11A2 exerts modulatory effects on STm infection via nutritional immunity. We used HCT116 colonic epithelial cells as an infection model to test this hypothesis *in cellulo*. Immunoblotting of whole-cell lysates with anti-SLC11A2 polyclonal antibodies indicated multiple variants are present in HCT116 cells, at ∼50 kDa and ∼80 kDa (Figure 3A). The SLC11A2 variants are due to alternative splicing which generates multiple mRNA isoforms (37, 70). STm infection did not affect the level of any SLC11A2 variants over a 12 h time course (Figure 3A; Figure S2A). Other highly abundant intestinal metal ion transporters - SLC40A1 (ferroportin 1, FPN) which exports iron and manganese from IECs (71, 72), SLC39A14 (ZIP14) which participates in manganese, iron, zinc and cadmium assimilation (7, 73–76), and CD71 which transports transferrin-bound iron (77) - were also unaffected over the infection timecourse (Figure S2B). As a positive control, we assessed levels of the iron-binding protein, lipocalin-2 (LCN2), in whole-cell lysates as STm infection enhances *LCN2* expression and LCN2 production in the intestinal mucosa in rhesus macaque ligated ileal loop and mouse colitis models *in vivo* (78). In agreement, the amount of cellular LCN2 significantly increased upon STm infection of HCT116 cells (Figure 3A, Figure S2A).

**Figure 3:**
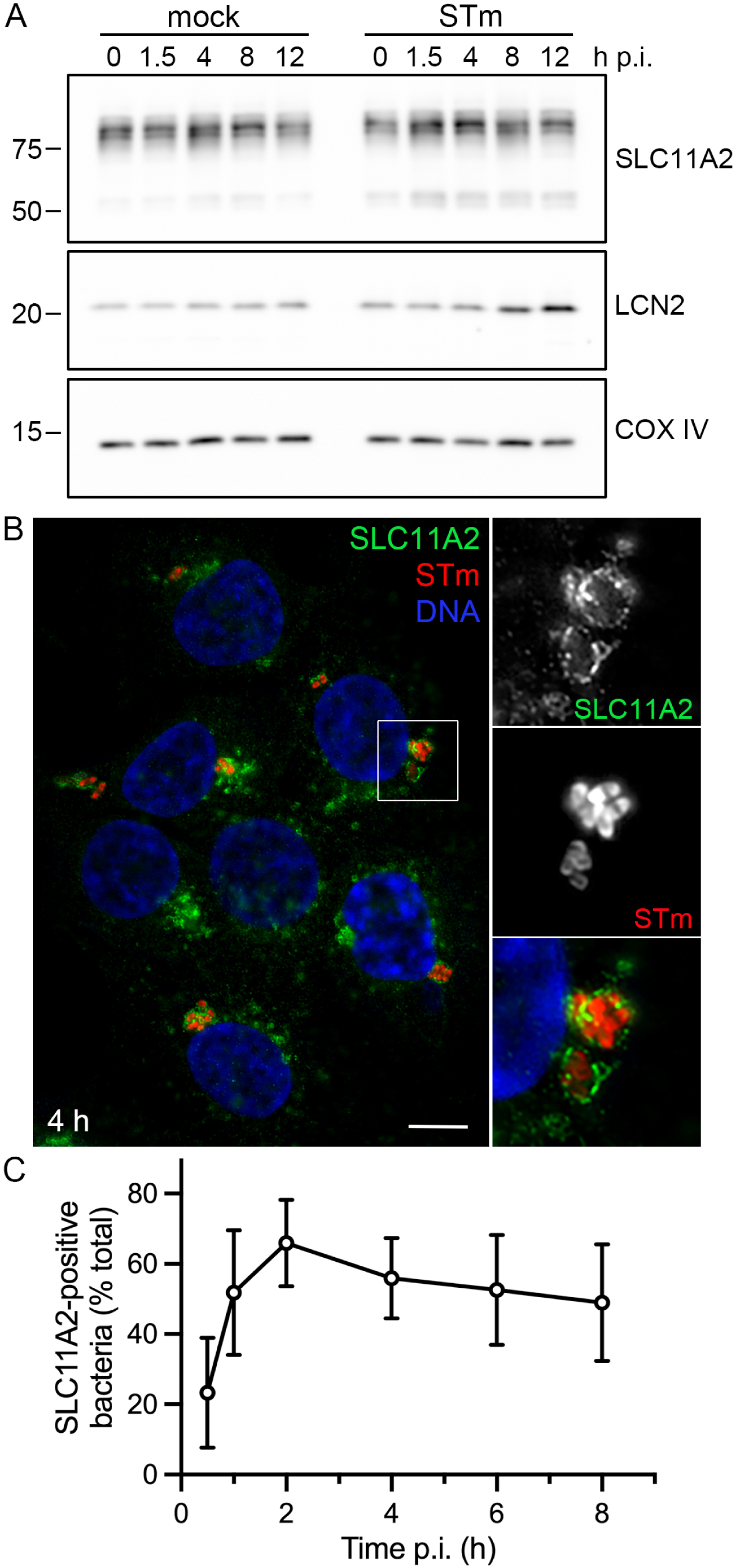
SLC11A2 is recruited to the *Salmonella*-containing vacuole. (A) HCT116 epithelial cells were mock-infected or infected with STm. Whole cell lysates were collected at the indicated times post-infection (p.i.). Proteins were separated by SDS-PAGE and subject to immunoblotting with antibodies against SLC11A2 (NRAMP2), lipocalin-2 (LCN2), a siderophore binding protein, and cytochrome c oxidase IV (COX IV). LCN2 is known to be induced following STm infection and is a positive control. COX IV is a loading control. Molecular mass markers are indicated on the left. Quantification of protein band intensity is shown in Figure S2. (B and C) HCT116 cells seeded on glass coverslips were infected with STm-mCherry and fixed with paraformaldehyde. Monolayers were permeabilized and immunostained with polyclonal antibodies against SLC11A2. DNA was stained with Hoecsht 33342. (B) Representative fluorescence microscopy image shows SLC11A2 recruitment (green) to the vacuolar membrane surrounding STm-mCherry (red) at 4 h p.i. Scale bar is 10 µm. (C) Timecourse of SLC11A2 recruitment. Samples were viewed by fluorescence microscopy and the number of SLC11A2-positive bacteria scored. Mean ± SD from n≥3 independent experiments.

SLC11A2 associates with the plasma membrane and endosomal network, particularly recycling endosomes, late endosomes and lysosomes, in various cell types (79–82). In uninfected HCT116 cells, SLC11A2 immunolocalized to numerous intracellular puncta concentrated in the perinuclear region (Figure 3B). Upon STm infection, SLC11A2 was temporally recruited to the *Salmonella*-containing vacuole (SCV) membrane (Figure 3B). Specifically, there was a progressive increase in SLC11A2 decoration of the SCV up to 2 h post-infection (p.i.) (66% SLC11A2-positive SCVs), followed by a plateau from 4-8 h p.i. (∼50% SLC11A2-positive SCVs) (Figure 3C). These acquisition kinetics are consistent with SCVs acquiring SLC11A2 as they mature along the endocytic pathway. In support of this idea, Slc11a2 is recruited to latex bead-containing phagosomes in mouse macrophages (79, 81).

To determine whether SLC11A2 exerts a protective effect against STm infection of IECs, HCT116 cells deficient in *SLC11A2* (SLC11A2 KO) were generated by CRISPR/Cas9 technology. Immunoblotting confirmed the loss of multiple SLC11A2 variants in SLC11A2 KO cells (Figure S3). SLC11A2 deletion did not have a compensatory effect on SLC40A1, SLC39A14 or CD71 protein levels, three other metal ion transporters in IECs (Figure S3). LCN2 levels were also unaffected by SLC11A2 inactivation (Figure S3). We compared the intracellular replication of STm in SLC11A2 wild type (WT) and KO cells using a gentamicin protection assay. Fold-replication was determined by dividing the colony forming units (CFUs) at 8 h or 16 h p.i. by those at 1 h p.i. There was a significant increase in bacterial replication in SLC11A2 KO cells compared to WT cells at 8 h (8.0±0.7-fold versus 6.6±1.1-fold, respectively; p<0.05, Student’s t-test) and 16 h p.i. (23.6±4.1-fold versus 12.1±3.9-fold, respectively; p<0.05, Student’s t-test) (Figure 4A). This increased proliferation was independently confirmed by scoring the number of STm-mCherry bacteria per cell by fluorescence microscopy. At 1 h and 4 h p.i., the mean number of bacteria per cell was comparable in WT and KO cells (Figure 4B). However, the mean number of bacteria per cell was increased in KO cells at 8 h p.i. and 12 h p.i., with statistical significance at the later timepoint (Figure 4B). STm occupies both vacuolar and cytosolic niches in epithelial cells (83). These two populations can be distinguished using a chloroquine (CHQ) resistance assay in conjunction with a gentamicin protection assay. Specifically, CHQ-resistant bacteria are cytosolic whereas CHQ-sensitive bacteria (total minus CHQ-resistant) are vacuolar (84, 85). We found that there was no significant difference in the number of vacuolar or cytosolic bacteria at 1.5 h p.i. when comparing SLC11A2 WT versus KO cells (Figure 4C). At 8 h p.i., cytosolic CFUs were equivalent but vacuolar CFUs were increased by 1.47-fold in SLC11A2 KO cells, albeit without statistical significance (p=0.14, Student’s t-test). By 16 h p.i., there was a significant increase in vacuolar (1.51-fold) and cytosolic (2.43-fold) populations in SLC11A2 KO cells (Figure 4C; p<0.05, Student’s t-test), indicating that STm replication in both intracellular niches is enhanced in the absence of SLC11A2.

**Figure 4:**
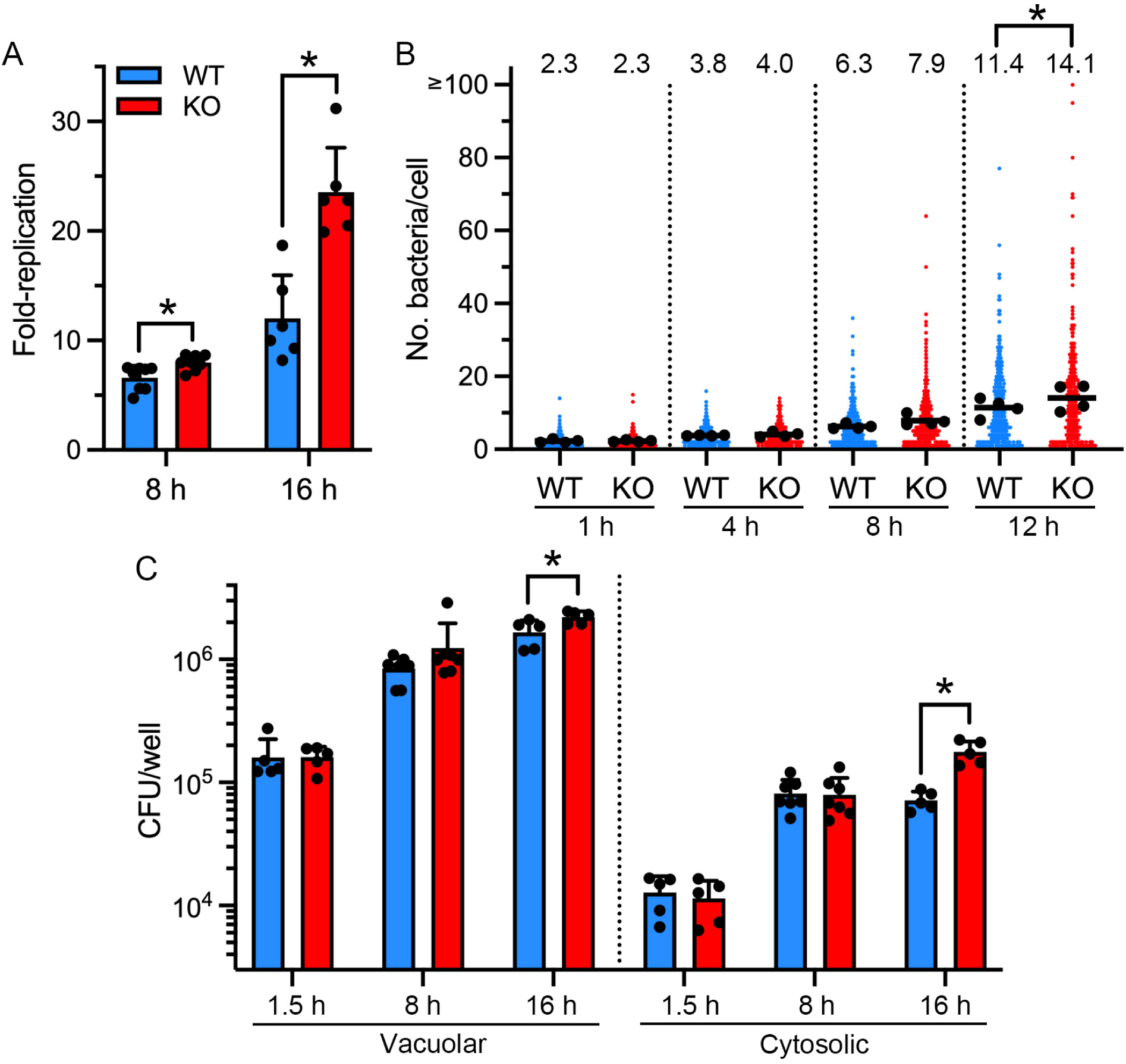
SLC11A2 limits *Salmonella* proliferation in intestinal epithelial cells. (A) HCT116 SLC11A2 WT and KO cells were infected with STm and the number of internalized bacteria at 1 h, 8 h and 16 h p.i. quantified by gentamicin protection assay. Fold-replication was determined by dividing the CFUs obtained for 8 h or 16 h p.i. by those at 1 h p.i. Mean ± SD from at least 6 independent experiments. *p<0.05, Student’s t-test. (B) HCT116 SLC11A2 WT and KO cells seeded on glass coverslips were infected with STm-mCherry. At 1 h, 4 h, 8 h and 12 h p.i., monolayers were fixed and the number of bacteria per cell was blindly scored by fluorescence microscopy. Small dots represent individual cells; large dots indicate the mean of each experiment; horizontal bars indicate the average of 4 independent experiments. *p<0.05, Student’s t-test. (C) HCT116 SLC11A2 WT and KO cells were infected with STm and the number of vacuolar and cytosolic bacteria at 1.5 h, 8 h and 16 h p.i. was determined by CHQ resistance assay in conjunction with a gentamicin protection assay. CHQ-resistant bacteria are cytosolic and CHQ-sensitive bacteria (total minus CHQ-resistant) are vacuolar. Mean ± SD from at least 5 independent experiments. *p<0.05, Student’s t-test.

### SLC11A2 withholds iron and manganese from intracellular STm

The presence of Slc11a1 alters the expression of SPI2-associated genes (*ssrA*, *sseA* and *sseJ*) but not SPI-1 associated genes (*hilA*) or *phoP* in mouse macrophages (86, 87). We did not detect any overt change in the temporal expression of SPI2-(*ssaG*) or SPI1-associated (*prgH*) genes in SLC11A2 WT or KO epithelial cells (Figure S4), discounting altered virulence gene expression as a likely reason for the increased bacterial replication. Instead, we hypothesized that SLC11A2 restricts intracellular STm replication via metal ion starvation. There is no clear consensus as to which metals SLC11A2 transports across the plasma membrane into mammalian cells. From transport studies with brush-border membrane vesicles, *Xenopus* oocytes and HEK293 cells, the substrate specificity of SLC11A2 is broad, *i.e.* Cd^2+^, Fe^2+^, Co^2+^, Mn^2+^, Ni^2+^, Pb^2+^, Cu^2+^ and Zn^2+^ (33, 37, 67–69). Therefore, we used an unbiased approach to compare metal ion levels in SLC11A2 WT and KO cells. Mock-infected and STm-infected HCT116 cells were collected at 8 h p.i. and metal ion concentrations (manganese, cadmium, iron, cobalt, nickel, zinc, copper and magnesium) were measured in cellular homogenates by inductively-coupled plasma mass spectrometry (ICP-MS). ^34^S was used as a reference element for normalization across samples (88). Cadmium and magnesium were the metals in lowest and highest abundance, respectively, in HCT116 cells (Figure S5). Comparing uninfected SLC11A2 WT and KO cells, cadmium ([^111^Cd]/[^34^S] of 1.28×10^−6^ ± 2.68×10^−7^ for WT, 9.86×10^−7^ ± 1.481×10^−7^ for KO; p=0.058, Student’s t-test) and iron levels ([^56^Fe]/[^34^S] of 2.11×10^−3^ ± 1.50×10^−4^ for WT, 1.91×10^−3^ ± 1.24×10^−4^ for KO; p=0.15, Student’s t-test) trended lower in SLC11A2 KO cells (Figure S5). The abundance of other metals – ^55^Mn, ^59^Co, ^60^Ni, ^66^Zn, ^63^Cu and ^24^Mg – was unchanged in SLC11A2 KO cells (Figure S5). STm infection in WT cells caused a significant reduction in cadmium ([^111^Cd]/[^34^S] of 1.28×10^−6^ ± 2.68×10^−7^ for mock, 8.46×10^−7^ ± 1.87×10^−7^ for infected; p<0.05, Student’s t-test) and magnesium ([^24^Mg]/[^34^S] of 0.154 ± 0.0017 for mock, 0.150 ± 0.00038 for infected; p<0.05, Student’s t-test) (Figure S5). For SLC11A2 KO cells, infection led to a significant increase in cobalt content (^59^Co/^34^S of 5.27×10^−6^ ± 4.98×10^−7^ for mock, 5.93×10^−6^ ± 3.14×10^−7^ for infected; p<0.05, Student’s t-test) but did not significantly alter the levels of other metals. Iron was the only metal found to be significantly different between WT and SLC11A2 KO cells upon STm infection ([^56^Fe]/[^34^S] of 2.08×10^−3^ ± 5.84×10^−5^ for WT, 1.94×10^−3^ ± 9.28×10^−6^ for KO; p<0.05, Student’s t-test) (Figure S5). Overall, our unbiased quantitative analysis hints that a subset of metals is altered in SLC11A2 KO cells compared to WT cells under basal and infection conditions.

To complement the population-based ICP-MS analysis, we used STm as a fluorescent biosensor to detect intracellular metal concentrations at the single-cell level. We previously established that P*iroN-gfpmut3* responds to iron >> cobalt, P*sitA*-*gfpmut3* to iron and manganese >> cobalt, P*zinT*-*gfpmut3* to zinc >> cobalt, and P*mgtC*-*gfpmut3* to magnesium (56). HCT116 SLC11A2 WT and KO cells were infected with STm-mCherry harboring these fluorescent reporters, monolayers were fixed at 4 h, 8 h and 12 h p.i. and the number of GFP-positive bacteria was scored by fluorescence microscopy. A general trend was that fewer STm-mCherry harboring P*iroN*-*gfpmut3* and P*sitA*-*gfpmut3* reporters were GFP-positive at 4 h, 8 h and 12 h p.i. in SLC11A2 KO cells (Figure 5A, S6A, S6C), but not for STm-mCherry harboring P*zinT*-*gfpmut3* or P*mgtC*-*gfpmut3* reporters (Figure 5A). We also considered heterogeneity in promoter-*gfp* expression in individual bacteria by quantifying the mean fluorescence intensity (MFI) per bacterium using ImageJ. At 8 h p.i., the average MFI for bacteria harboring P*iroN*-*gfpmut3* and P*sitA*-*gfpmut3* reporters was significantly higher in WT cells than SLC11A2 KO cells (1.77-fold for P*iroN*, 1.86-fold for P*sitA*, p<0.05, Student’s t-test; Figure 5B, 5C). A similar trend was observed at 4 h and 12 h p.i. (Figure S6B, S6D). By contrast, the average MFI for bacteria harboring P*zinT*-*gfpmut3* and P*mgtC*-*gfpmut3* reporters was unchanged in HCT116 WT and KO cells at 8 h p.i. (Figure 5B). The increased expression of *iroN* and *sitA* reporters at the population and single-cell levels, together with the ICP-MS analysis, indicates that SLC11A2 limits iron and manganese from STm during infection. Additionally, the comparable expression of *zinT* or *mgtC* reporters shows that the accessibility of zinc and magnesium to STm is equivalent in SLC11A2 WT and KO cells.

**Figure 5:**
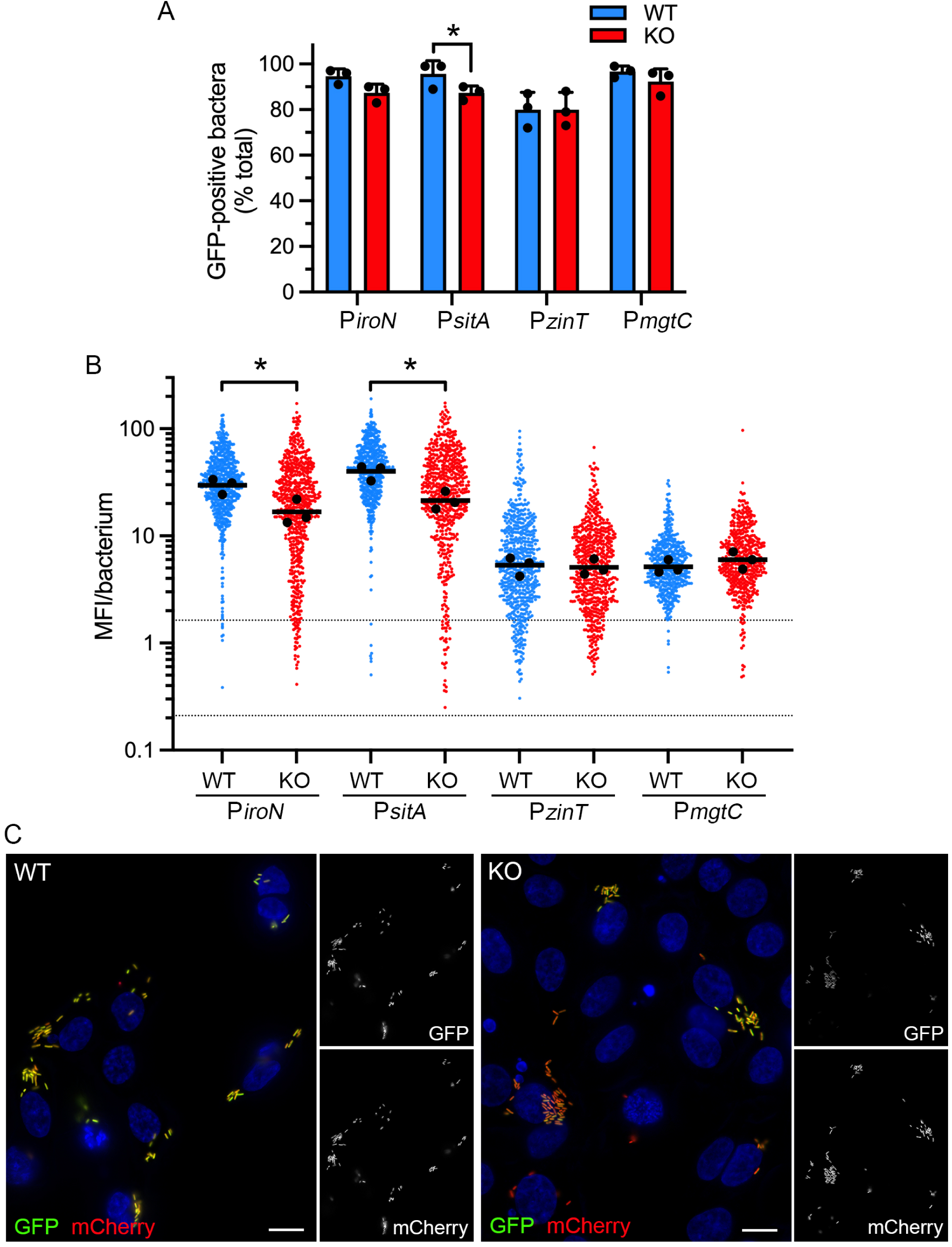
SLC11A2 withholds iron and manganese from intracellular *Salmonella*. (A) HCT116 SLC11A2 WT and KO cells seeded on glass coverslips were infected with STm-mCherry harboring the following fluorescent transcriptional reporters: P*iroN*-*gfpmut3*, P*sitA*-*gfpmut3*, P*zinT*-*gfpmut3*, P*mgtC*-*gfpmut3*. At 8 h p.i., monolayers were fixed and stained with Hoechst 33342 to label DNA. The number of GFP-positive bacteria was scored by fluorescence microscopy. Mean ± SD from 3 independent experiments. *p<0.05, Student’s t-test. (B) Quantification of the MFI of GFP signal by fluorescence microscopy and ImageJ analysis. Acquisition parameters (exposure time and gain) were the same for all reporters. Small dots represent individual bacteria; large dots indicate the mean of each experiment; horizontal bars indicate the average of three independent experiments. The dashed lines indicate the range of background fluorescence in the GFP channel for mCherry-STm (no reporter plasmid). *p<0.05, Student’s t-test. (C) Representative images of HCT116 SLC11A2 WT and KO cells infected with STm-mCherry P*sitA*-*gfpmut3* bacteria at 8 h p.i. Scale bars are 10 µm.

### MntH and EntC compete against SLC11A2 for trace metals

We next investigated whether bacteria deficient in metal acquisition were attenuated for growth in IECs, and if so, whether deletion of SLC11A2 would rescue this attenuation. HCT116 WT cells were infected with STm wild-type or mutants defective in high affinity Mn^2+^/low affinity Fe^2+^ import (Δ*sitA*Δ*mntH*), high affinity Zn^2+^/low affinity Mn^2+^ and Co^2+^ import (Δ*zupT*Δ*znuA*), Mg^2+^ import (Δ*mgtA*Δ*mgtB*) (24) or siderophore synthesis (Δ*entC*). Growth over 16 h was quantified by gentamicin protection assay. Compared to wild-type bacteria (15.9±4.6-fold), replication of Δ*entC* (10.4±1.4-fold) and Δ*mntH*Δ*sitA* (6.5±1.5-fold) bacteria was significantly decreased in SLC11A2 WT cells (Figure 6A). Compared to wild-type bacteria, Δ*entC* bacteria had decreased replication in the SCV, whereas Δ*mntH*Δ*sitA* bacteria were replication-deficient in both the SCV and cytosol (Figure 6B, 6C). Plasmid-borne complementation with *entC* and *mntH* (but not *sitA*) rescued the replication defect of the respective deletion mutants in SLC11A2 WT cells (Figure 6D, 6E). The lack of complementation by plasmid-borne *sitA* suggests that the SLC11A2-dependent restriction of Δ*mntH*Δ*sitA* bacteria is primarily due to the loss of MntH. This result might be explained by the different pH optima of MntH and SitABCD transport capacities; in acidic conditions (such as those found in the SCV lumen), MntH uptake of Mn^2+^ is maximal whereas SitABCD is largely inactive (60). Notably, fold-replication for Δ*zupT*Δ*znuA* and Δ*mgtA*Δ*mgtB* mutants was comparable to wild-type bacteria in SLC11A2 WT cells (Figure 6A). In SLC11A2 KO cells, fold-replication of all the deletion strains was indistinguishable from wild-type bacteria (Figure 6A). The Δ*zupT*Δ*znuA* mutant replicated better than wild-type bacteria in the cytosol of WT and KO cells (Figure 6C), indicating that this phenotype is SLC11A2-independent. We reason that deletion of SLC11A2 rescues the replication defect of Δ*entC* and Δ*mntH*Δ*sitA* bacteria because it removes the host-advantage in the competition for iron and manganese. Overall, these results directly connect the bioavailability of these two metals with SLC11A2-mediated restriction of STm growth in IECs.

**Figure 6:**
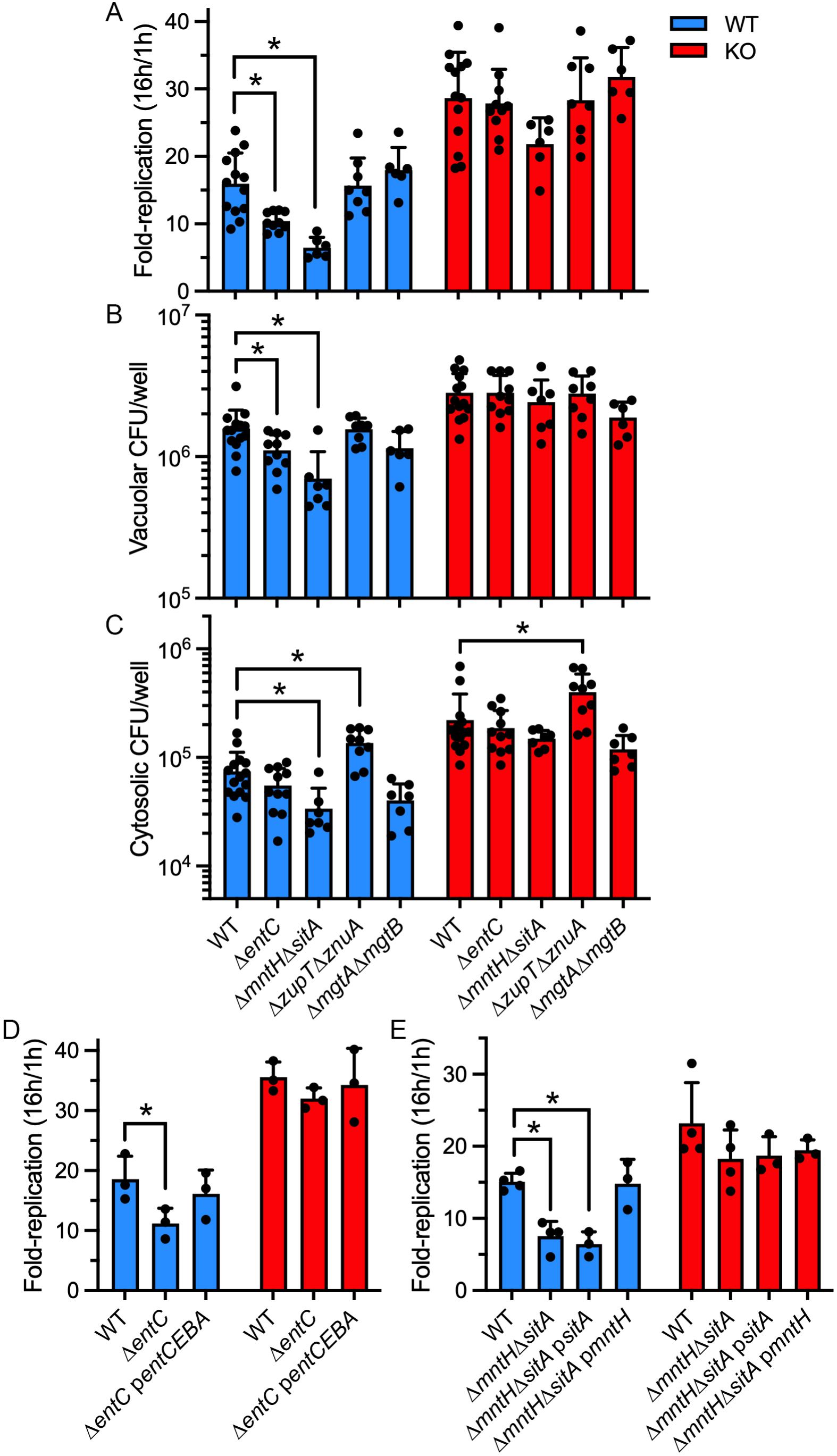
Iron and manganese acquisition by *Salmonella* counter SLC11A2 activity. (A) HCT116 SLC11A2 WT and KO cells were infected with STm wild-type (WT), Δ*entC*, Δ*mntH*Δ*sitA*, Δ*zupT*Δ*znuA* or Δ*mgtA*Δ*mgtB* bacteria. Fold-replication was determined by gentamicin protection assay by dividing CFUs at 16 h by those at 1 h p.i. (B and C) The number of vacuolar (B) and cytosolic (C) bacteria at 16 h p.i. was determined by CHQ resistance assay in conjunction with a gentamicin assay. CHQ-resistant bacteria are cytosolic whereas CHQ-sensitive bacteria (total minus CHQ-resistant) are vacuolar. Mean ± SD from ≥6 independent experiments. *p<0.05, ANOVA with Dunnett’s post-hoc test. (D and E) *In trans* complementation. (D) HCT116 SLC11A2 WT and KO cells were infected with STm wild-type, Δ*entC* and Δ*entC* pWSK29-*entCEBA* (p*entCEBA*) bacteria or (E) STm wild-type, Δ*mntH*Δ*sitA*, Δ*mntH*Δ*sitA* pWSK29-*sitA* (p*sitA*) and Δ*mntH*Δ*sitA* pWSK29-*mntH* (p*mntH*) bacteria. Fold-replication was determined by gentamicin protection assay by dividing CFUs at 16 h by those at 1 h p.i. Mean ± SD from at 3-4 independent experiments. *p<0.05, ANOVA with Dunnett’s post-hoc test.

## Discussion

STm face stiff competition for metal ions whether extracellular and intracellular, and at both intestinal and extra-intestinal sites. Studies on metal ion competition between STm and its mammalian host have largely used C57BL/6 (*Slc11a1*^G169D^ *i.e. Slc11a1* null background) mice and streptomycin pretreatment to alter the resident gut microbiota. In this mouse colitis model, STm efficiently colonizes the cecum (although most bacteria remain extracellular (89)), and elicits severe neutrophilic inflammation and mild secretory responses in the cecum and proximal colon, then systemically spreads to the liver and spleen ∼48 h after inoculation (90). This infection model has been instrumental in defining much of what we know about the cell- and tissue-specificity of nutritional immunity against *S. enterica*. For example, in response to STm infection, neutrophils and other inflammatory cells release calprotectin, a metal chelator (91), leading to increased fecal calprotectin (a proxy for lumenal content), and reduced fecal zinc and manganese (92). Likewise, increased Lcn2 production in the intestinal mucosa and secretion into the lumen promotes the sequestration of enterobactin produced by STm and other enteric bacteria (90). A small peptide hormone, hepcidin, is also released from hepatocytes to deplete extracellular iron (86–88). As a result, serum and fecal iron levels decline markedly in STm-infected mice (92, 94, 95). Additionally, decreased intracellular ferric iron has been reported in duodenal enterocytes during long-term STm infection of Sv129S6 (*Slc11a1*^+/+^) mice (92). Here we show that, in addition to iron, intracellular STm experience manganese and zinc limitation in the bovine intestinal mucosa (Figure 2).

The SLC11 transporters appear to play site- and cell-specific roles during infection. *Slc11a1* expression is typically ∼50-fold higher than that of *Slc11a2* in phagocytes (79), suggesting that Slc11a1 functions dominate in this cell type. In concordance, Slc11a1 limits STm replication in the spleen, liver and mesenteric lymph nodes of mice (24, 48, 49), organs in which STm primarily resides within macrophages, but has little to no effect on STm replication in the mouse gut (48, 49). Mice deficient in *Slc11a2* (*Slc11a2*^-/-^) die within 7 days of birth (38), which has limited studies on its contribution to host defense. *Slc11a2*^-/-^ bone marrow-derived macrophages support increased bacterial burdens (96) with the caveat that this study used macrophages derived from C57BL/6 mice, which have a non-functional Slc11a1 (*Slc11a1*^G169D^). Based on our results, we propose that SLC11A2 promotes host defense in the gut epithelium via iron and manganese sequestration. Upon STm infection of chickens, there is a robust increase in *Slc11a1* expression in the liver and spleen 1-3 days p.i. and the cecum by day 7 p.i. (97). *Slc11a2* expression in the liver and cecum peaks later, at day 9 p.i. (97), suggesting a temporal, tissue-specific defensive role for SLC11 transporters in avian hosts as well.

*S. enterica* employs numerous strategies to fight back in the war for metals. With regard to iron, STm produces a second siderophore, salmochelin, a glucosylated version of enterobactin that is not bound by LCN2 (98). An *iroN* (receptor for ferric-bound enterobactin and salmochelin) deletion mutant is defective for cecal colonization in streptomycin-pretreated mice at 48 h p.i. but not in the absence of intestinal inflammation (78). In a transposon screen for STm mutants defective for intestinal colonization in calves and chickens at day 4 p.i., and pigs at day 3 p.i., enterobactin and salmochelin synthesis mutants showed variable degrees of fitness depending on the Tn7 insertion site (99). Our study of STm and metal limitation is focused on intestinal tissue and cellular colonization, which distinguishes it from other studies which largely interrogate STm fitness in lumenal contents (78, 91, 100, 101). Intracellular STm experience iron deprivation in the intestinal mucosa (Figure 2), specifically within IECs and lamina propria phagocytes, and deploy siderophores to compete with SLC11A2 for iron and manganese in gut epithelial cells (Figure 6). Despite this iron stress, an Δ*entC* mutant does not have a colonization defect in the intestinal lumen or mucosa of a bovine host at 2 h (Figure S1) or 8 h (Figure 2), in agreement with an earlier study of various iron acquisition mutants (*iroN*; *tonB*, Fe^3+^ uptake; *feoB*, Fe^2+^ uptake) in bovine ligated loops (102). STm infection of ligated intestinal loops models the very early interactions between bacteria and the intestinal mucosa while avoiding bottlenecks associated with traversal of the stomach, duodenum, and proximal jejunum. An intrinsic limitation of this infection model is the limited number of bacterial doublings in the intestinal lumen over the short duration of the experiment (typically 8 h). STm deletion mutants are grown in rich media prior to loop inoculation, and do not have a growth defect in this media (56, 103, 104). It is possible that bacteria are sufficiently metal replete to allow them to survive or/and undergo limited replication in the face of the host inflammatory response in intestinal loops. Additionally, STm siderophore synthesis mutants can still acquire iron through other mechanisms *in vivo* such as ferric-bound xenosiderophores produced by resident microbiota (105, 106). Despite these caveats, it is clear that siderophore-mediated iron acquisition contributes to STm fitness at systemic sites (107), and at later times in the inflamed intestine (78) when high-affinity iron binding proteins such as LCN2 and hepcidin have been released, and bacteria are replicating (90).

STm can circumvent calprotectin-mediated metal limitation in the mouse gut through the expression of a high-affinity Zn^2+^ importer, ZnuABC (91), and two high-affinity Mn^2+^/low-affinity Fe^2+^ importers, MntH and SitA (61). Here, we identified transcriptional signatures of zinc, manganese and iron limitation in bovine intestinal mucosa and that MntH counters SLC11A2-mediated manganese restriction within IECs. Surprisingly, STm mutants defective for Mn^2+^ transport (Δ*mntH*, Δ*sitA*, Δ*mntH*Δ*sitA*) colonize the mouse cecum at levels equivalent to wild-type bacteria at 96 h p.i. (61). In the bovine intestine, we also saw no difference in lumenal or tissue colonization between wild-type and Δ*mntH*Δ*sitA* bacteria at 2 h (Figure S1) or 8 h (Figure 2C). The lack of an overt phenotype might be explained by Δ*mntH*Δ*sitA* mutants being able to acquire trace amounts of Mn^2+^ via ZupT and other unidentified mechanisms (108, 109) and/or the metal-replete growth conditions prior to loop inoculation. In mixed infections, wild-type bacteria outcompete *mntH*, *sitA* and *mntH sitA* deletion mutants in the small intestine and spleen of C57BL/6, BALB/C (*i.e.*, *Slc11a1*^-/-^) and C3H/HeN (i.e., *Slc11a1*^+/+^) mice (100, 110), indicating that Mn^2+^ acquisition does provide STm with a competitive advantage at intestinal and extra-intestinal sites. Distinct from Mn^2+^ uptake mutants, a Zn^2+^ uptake mutant is severely compromised in the mouse colitis model; cecal colonization by a Δ*znuA* mutant is reduced ∼200-fold at 96 h p.i. (100). However, neither the ZnuABC transporter nor the IroN receptor provide a growth advantage in the mouse in the absence of gut inflammation (100), or in the bovine loop model (Figure 2) (102).

We observed phenotypic heterogeneity in STm metal limitation *in cellulo* and *in vivo*. Activation of *iroN*, *sitA*, and *zinT* promoters was observed in the majority of STm as early as 4 h p.i. *in cellulo* (Figure S6) but the variation in transcriptional activity (*i.e*., GFP intensity) between individual bacteria was great (Figure S6, Figure 5), even within the same IEC (Figure 5). By contrast, the minority of bacteria experience metal limitation within both epithelial cells and cells of the lamina propria *in vivo* (Figure 2). This could be due to fewer intracellular bacteria and proportion of infected host cells *in vivo* compared to *in cellulo*, which would lead to a slower exhaustion of the available pool of metals. We also observed inter-bacterial variations in *iroN*, *sitA* and *zinT* and promoter activity at the tissue and cellular level *in vivo* (Figure 2). These data imply that metal limitation is not evenly distributed within the gut tissue or tissue microcompartment, consistent with reports of differential iron availability in *Staphylococcus aureus* infected tissues (111, 112). Given not all bacteria experience the same degree of metal limitation during infection, culturing bacteria from homogenized tissue may mask subtle phenotypes like differential fitness within individual cells or tissue microcompartments.

While the phagocyte-specific transporter Slc11a1/SLC11A1 has assumed a place of prominence in nutritional immunity, here we show that SLC11A2 in IECs also shapes infection outcome via metal deprivation. Under basal conditions, SLC11A2 localizes to the plasma membrane and intracellular vesicles in IECs (79–82). Upon STm infection, SLC11A2 is temporally recruited to SCVs (Figure 3). In *SLC11A2* KO IECs, we observed enhanced STm proliferation in both the SCV and cytosol (Figure 4) and decreased *iroN* and *sitA* promoter activity, but not *zinT* or *mgtC* promoter activity (Figure 5). Collectively, our results support a scenario whereby SLC11A2 transports iron and manganese out of the phagosome to deprive vacuole-resident bacteria of these metals. Vacuolar STm employ MntH and EntC to compete against SLC11A2-driven efflux of these metals. How might SLC11A2 limit the cytosolic population of STm in IECs? Iron and manganese homeostasis are intimately associated with oxidative stress (113, 114) and we have previously shown that STm are exposed to oxidative stress while in the cytosol (56). One possible explanation is metal intoxication, *i.e.*, SLC11A2 imports metals into the cytosol, induces metal toxicity-related oxidative stress and thereby restricts cytosolic STm. SLC11A2 is one of 32 metal ion transporters from six SLC families - SLC11, SLC30, SLC31, SLC39, SLC40 and SLC41 - in mammals (115, 116). Our study unequivocally shows that SLC11A2-mediated sequestration of trace metals is an important IEC defense measure against STm, but whether these other SLC family members play a similar role in nutritional immunity in the gastrointestinal tract remains an unanswered question, one worthy of attention.

## Materials and Methods

### Bacterial strains and plasmids

*S.* Typhimurium (STm) ST4/74 was the wild-type (WT) strain used in this study (117, 118). WT bacteria constitutively expressing codon-optimized *mCherry* on the chromosome (*glmS::Ptrc-mCherryST::*FRT), STm-mCherry, were created by P22 transduction from STm SL1344 WT *glmS::Ptrc-mCherryST*::Cm (119). The following mutants were created by transduction using P22 lysates prepared from the STm 14028s single gene deletion mutant library (120): Δ*entC*::Kan, Δ*znuA*::Kan, Δ*mntH*::Kan Δ*sitA*::Cm, Δ*zupT*::FRT Δ*znuA*::Kan, Δ*mgtA*::Kan Δ*mgtB*::Cm and Δ*invA*::FRT Δ*ssaD*::Kan. pCP20 (121) was used for Flp-mediated excision of antibiotic resistance gene cassettes. The transcriptional reporters pMPMA3ΔPlac-P*iroN*-*gfpmut3.1,* pMPMA3ΔPlac-P*sitA*-*gfpmut3.1,* pMPMA3ΔPlac-P*zinT*-*gfpmut3.1*, pMPMA3ΔPlac-P*mgtC*-*gfpmut3.1*, pMPMA3ΔPlac-P*prgH*-*gfp(LVA)*, and pMPMA3ΔPlac-P*ssaG*-*gfp(LVA)* have been described previously (56, 122) and were electroporated into STm-mCherry bacteria. The transcriptional fusion plasmids were stably maintained by bacteria over the duration of our experiments *in cellulo* and *in vivo* (not shown). For plasmid-borne complementation, the coding sequences and upstream regulatory regions of *sitA* and *mntH* were cloned into the low copy number plasmid, pWSK29 (123). Oligonucleotides are listed in Table S1. The pWSK29-*entCEBA* plasmid has been described previously (56).

### Mammalian cell culture

*SLC11A2* wild type (WT) and knockout (KO) HCT116 (human colorectal carcinoma epithelial cells) cell pools were generated using CRISPR/Cas9 technology by Synthego. For the WT cell pool, parental cells were electroporated with SpCas9 only and confirmed to be unedited at the *SLC11A2* locus. For the KO cell pool, parental cells were electroporated with SpCas9 and three *SLC11A2*-specific sgRNAs targeting exon 5 (CUAGACUGGGAGUGGUUACU, GUUGCUCUGGAUCCUUCUGU, GACAAUACAUUGCUCACCUU). Gene editing efficiency was >99.9%. Cells were maintained at 37°C and 5% CO_2_ in McCoy’s 5A (Iwakata and Grace modification, Gibco) medium containing 10% (v/v) heat-inactivated fetal calf serum (FCS; Gibco) and used within 15 passages of receipt. SLC11A2 was not detected in *SLC11A2* KO cell lysates by immunoblotting up to passage 15 (not shown).

### Bacterial infection of mammalian cells

HCT116 *SLC11A2* WT and KO cells were seeded 48 h prior to infection at a density of 8×10^4^ cells/well in collagen-coated (rat tail collagen I, Corning) 24-well plates or 3.2×10^5^ cells/well in 6-well plates (Thermo Scientific Nunc). Growth media was replaced with McCoy’s 5A media containing 1% (v/v) heat-inactivated FCS 20-22 h prior to infection. Bacterial cultures were grown overnight (16-20 h) in 2 ml LB-Miller broth (BD Difco) with shaking (220 rpm) at 37°C, then sub-cultured 1:33 into 10 ml LB-Miller broth in 125 ml Erlenmeyer flasks. Growth continued at 37°C for 3.5 h with shaking (220 rpm). Cell monolayers were infected with bacterial subcultures (MOI ∼100) for 10 minutes. Gentamicin protection assays and CHQ resistance assays (400 µM CHQ) were as described previously (85), except that McCoy’s media containing 1% FCS was used for all infections. Monolayers were solubilized in 0.2% (w/v) sodium deoxycholate (Sigma) and serial dilutions were plated on LB-Miller agar (BD Difco) for enumeration of colony forming units (CFUs).

### Total Metals Quantitation

*SLC11A2* WT and KO cells were seeded in 4×15 cm tissue culture treated dishes (Nunc) at 1×10^7^ cells per dish. Growth media was changed to McCoy’s containing 1% FCS the next day, then mock-infected or STm-infected for 8 h the following day. Monolayers were washed with 30 ml tissue culture grade phosphate buffered saline without calcium and magnesium (PBS^--^, Corning), then scraped into 10 ml PBS^--^, collected and pooled in a metal-free 50 ml centrifuge tube (VWR) and centrifuged at 400xg for 5 min. The supernatant was removed, the cell pellet resuspended in 10 ml PBS^--^, then transferred to a metal-free 15 ml centrifuge tube (VWR) and centrifuged again. The supernatant was removed and the wet weight of the cell pellet recorded. Samples were digested in 600 µL 70% Optima-grade nitric acid at 65°C overnight, then diluted with UltraPure water to 20% nitric acid for analysis. Elemental quantification was conducted using an Agilent 7700 ICP-MS attached to an ASX-560 autosampler. The settings for analysis were cell entrance = −40 V, cell exit = −60 V, plate bias = −60 V, OctP bias = −18 V, and helium flow = 4.5 ml/min. Optimal voltages for extract 2, omega bias, omega lens, OctP RF, and deflect were empirically determined. Calibration curves for elements were generated using ARISTAR ICP standard mix. Samples were introduced by peristaltic pump with 0.5-mm-internal-diameter tubing through a MicroMist borosilicate glass nebulizer. They were initially taken up at 0.5 rps for 30 seconds, followed by 30 seconds at 0.1 rps to stabilize the signal. Spectrum mode analysis was performed at 0.1 rps, collecting three points across each peak and conducting three replicates of 100 sweeps for each element. The sampling probe and tubing were rinsed with 2% nitric acid for 30 seconds at 0.5 rps between each sample. Data were acquired and analyzed using Agilent MassHunter workstation software version A.01.02. The concentration of each metal (in ppb) was normalized to that of ^34^S (in ppb) in each sample.

### Bovine infections

The Institutional Animal Care and Use Committee of University of Wisconsin-Madison approved the bovine infection experiment (Protocol number V006249). All experiments were performed in accordance with the PHS “Guide for the Care and Use of Laboratory Animals” in AAALAC-approved animal facilities. Holstein cross-bred calves were obtained from a University of Wisconsin-Madison farm herd. Genetic variations in the bovine *SLC11A1* gene have been described but none are known to negatively affect the expression or targeting (124). Calves were separated from the dam and administered colostrum on the farm and transferred to AAALAC-approved large animal housing facilities within 5 days of birth with housing in individual or grouped isolation rooms. Additional colostrum replacer was administered if determined necessary by measurement of serum total protein to estimate adequate passive transfer of immunity. Calves were fed milk replacer at 10-20% body weight per day with free choice access to water, hay, and calf starter. Selective fecal cultures were performed at least twice weekly and all calves had at least one negative fecal culture for *Salmonella* prior to surgery. For calves with *Salmonella* isolates obtained from feces one isolate was serotyped at the Wisconsin Veterinary Diagnostic Laboratory and no calf was harboring any *Salmonella* serotype B.

In preparation for ligated loop infections, bacteria were grown overnight at 37°C with shaking (225 rpm) in LB broth. Overnight cultures were sub-cultured 1:100 into LB broth and grown for approximately 4 hours at 37°C with shaking (225 rpm). Bacteria were washed twice in PBS and cell concentration was normalized by optical density (OD_600_). Actual inoculum dose was determined by serial dilution and plating.

At 3-6 weeks of age, calves were placed under general anesthesia with intravenous propofol and maintained with isoflurane inhalant for ligated jejuno-ileal ligated loop surgery as previously described with minor modifications (66). Briefly, calves were positioned in left lateral recumbency and a right flank incision was made. Up to thirty-eight 3- to 6-cm loops were tied in the ileum and terminal jejunum with 1-cm spacers between adjacent loops. Loop lengths were recorded prior to inoculation of 2 ml PBS with ∼10^9^ CFU of the indicated bacterial strains. Infected intestinal segments were returned to the abdomen, the incision closed, and the calves were maintained under inhalant anesthesia for the duration of the experiment. Calves were euthanized by intravenous pentobarbital after 2 h or 8 h incubation of infected loops. After euthanasia, the incision was opened, and all loops were excised individually. Following excision, intestinal fluid and tissue samples were harvested and processed. Fluid volume was calculated by excising individual loops and weighing escaped lumenal fluid on a sterile Petri dish. Lumenal fluid was then transferred to 1 mL PBS to allow for bacterial enumeration. Intestinal tissues were first processed for tissue fixation samples to maintain tissue integrity with the remaining tissue sample washed twice in PBS to remove ingesta and non-adherent bacteria. Washed tissues were then cut in half with one segment processed to quantify tissue-associated bacteria, and the other half was treated with gentamicin (50 µg/mL) for 30 minutes at 37°C to quantify intracellular bacteria. After gentamicin treatment, tissues were washed twice with PBS to remove remaining gentamicin. Samples were homogenized, serially diluted, and plated on the appropriate antibiotics for CFU enumeration.

### Immunostaining and fluorescence microscopy

*SLC11A2* WT and KO cells were seeded on acid-washed, collagen-coated 12 mm glass coverslips (#1.5 thickness, Fisher Scientific) in 24-well tissue culture plates. For HCT116 cells infected with STm-mCherry harboring transcriptional reporters, monolayers were washed once in PBS, then fixed with 2.5% (w/v) paraformaldehyde (PFA) in PBS for 10 min at 37°C. DNA was stained with Hoechst 33342 (Invitrogen) and coverslips were mounted onto glass slides using Mowiol. For immunodetection of SLC11A2, monolayers were infected with STm-mCherry as described above, washed once in PBS, then fixed in 2.5% PFA for 10 min at 37°C. Cell monolayers were blocked/permeabilized in 10% (v/v) normal goat serum (Gibco)/0.2% (w/v) saponin in PBS (blocking buffer) for 20 min at room temperature (RT), then incubated with rabbit monoclonal anti-SLC11A2/DMT1 (D3V8G, Cell Signaling) diluted 1:100 in blocking buffer for 30-45 min at RT. Coverslips were washed three times with PBS and incubated for a further 30–45 min at RT with goat anti-rabbit Alexa Fluor 488 antibodies (1:400 dilution; Life Technologies) in blocking buffer. After three washes in PBS, cells were incubated with Hoechst 33342 (1:10,000 dilution; Invitrogen) for 1 min before mounting in Mowiol on glass slides. Samples were cured overnight at RT.

Samples were visualized on a Leica Thunder DM4 upright fluorescence microscope and images were scored for number of bacteria per cell, proportion of GFP-positive bacteria or SLC11A2-positive bacteria, and quantification of GFP fluorescence. The number of bacteria in 100 infected cells across ≥5 random fields of view in the mCherry channel, as well as the corresponding number of GFP-positive bacteria, were manually scored at 100x magnification. To minimize the impact of observer bias (125), samples were blinded prior to analysis.

For quantification of the mean fluorescence intensity (MFI) of GFP signal in individual bacteria, image acquisition parameters were determined using bacteria harboring P*sitA*-*gfp*, the transcriptional reporter with the highest GFP intensity, and were applied to all transcriptional reporters for that timepoint. ImageJ software was used to quantify the activity of transcriptional reporters at the individual bacterium level as previously described (56). Sample identity was blinded to the observer. From grayscale images of ≥5 randomly selected fields of view on the mCherry channel (representative of all bacteria), 1-5 well-defined bacteria per infected cell were arbitrarily chosen, manually outlined, and converted into a binary image. The MFI of each bacterium was then determined by transferring the mask from the mCherry channel to the corresponding GFP channel image. The number of GFP-positive bacteria and total number of bacteria were quantified manually from each image.

Bovine intestinal tissue fixation and staining was as described previously with minor modifications (126). Bovine tissue samples were transferred to tissue cassettes and placed into 10% buffered formalin for 24 h. Samples were then floated into 20% (w/v) sucrose with 0.05% (w/v) sodium azide and stored at 4°C until use. Tissue samples were submitted to the University of Wisconsin-Madison Translational Research Initiatives in Pathology (TRIP) laboratory for tissue embedding, freezing, and tissue sectioning (10 µm). For tissues infected with STm-mCherry harboring GFP transcriptional reporters, sections were rehydrated in PBS for 5 min, blocked with 2% normal donkey serum (EMD Millipore), 1% bovine serum albumin (BSA, CELLect, MP Biomedicals), 0.1% Triton X-100 (MP Biomedicals), and 0.05% Tween 20 (Calbiochem) in PBS (NDS/TX100/Tween/PBS) for 30 min at RT. For immunostaining, sections were rehydrated with PBS for 5 min, followed by permeabilization and blocking with NDS/TX100/Tween/PBS for 45 min at RT. Sections were incubated overnight at 4°C in a humidified chamber with rabbit monoclonal anti-SLC11A2 (1:100 dilution, D3V8G, Cell Signaling Technology) or rabbit polyclonal anti-human SLC11A1 (1:100 dilution, NBP1-87809, NovusBio) (antibody was developed against a human SLC11A1 peptide sequence that shares 87% identity with bovine SLC11A1) in 1% BSA, 0.1% TX100, and 0.05% Tween 20 in PBS, followed by 3 washes for 5 min each in 0.05% Tween 20 in PBS (PBST). Donkey anti-rabbit Alexa Fluor 488 and phalloidin Alexa Fluor 647 (Invitrogen) were diluted 1:400 in 0.1% Triton X-100 in 0.05% Tween 20 in PBS and incubated for 1 h at RT. Sections were washed 3 times for 5 min each in PBST, then covered with ProLong Gold antifade reagent with 4’,6-diamidino-2-phenylindole (DAPI; Life Technologies), followed by a glass coverslip (Corning). Samples were cured overnight at RT. Slides were viewed at 63x or 100x on a Leica Thunder DM4 upright fluorescence microscope. Images were processed using Adobe Photoshop 2024.

### Immunoblotting

Monolayers in 6-well plates were washed twice with PBS prior to lysis in boiling 1.5x SDS-PAGE sample buffer. Proteins were separated by SDS-PAGE and subsequently transferred to 0.2 µm nitrocellulose membranes (Amersham). Membranes were blocked at room temperature for 1-2 h with Tris-buffered saline (TBS) containing 5% (w/v) skim milk powder and 0.1% (v/v) Tween-20 (TBST-milk) for SLC11A2, SLC40A1, CD71 and COX IV, or TBST with 3% (w/v) BSA (Millipore) (TBST-BSA) for LCN2 and SLC39A14. Membranes were then incubated overnight at 4°C with the following primary antibodies in TSBT-milk or TBST-BSA (LCN2, SLC39A14): rabbit monoclonal anti-SLC11A2 (D3V8G, 1:1,000 dilution, Cell Signaling), rabbit polyclonal anti-FPN1 (1:4,000 dilution, Proteintech), rabbit monoclonal anti-CD71 (D7G9X, 1:5,000 dilution, Cell Signaling), rabbit monoclonal anti-LCN2 (D4M8L, 1:1,000 dilution, Cell Signaling), rabbit monoclonal anti-SLC39A14/ZIP14 (E3H7D, 1:1000 dilution, Cell Signaling) or rabbit monoclonal anti-COX IV (3E11, 1:5,000 dilution, Cell Signaling). Membranes were then incubated with anti-rabbit IgG horseradish peroxidase (HRP)-conjugated secondary antibodies (1:10,000 dilution, Cell Signaling) in TBST-milk for 1 h at room temperature, followed by Supersignal West Femto Max Sensitivity ECL Substrate (Thermo). Chemiluminescence was detected using a GE Healthcare AI600 imager. ImageJ software was used to quantify protein band intensity relative to the COX IV loading control.

### Statistical analysis

All experiments were conducted on at least three separate occasions, unless otherwise indicated. Results are presented as mean ± SD, except for MFI data, where the geometric mean is presented. Statistical analyses were performed using: (i) Student’s t-test, (ii) one-way analysis of variance (ANOVA) with Dunnett’s post-hoc test, or (iii) two-way ANOVA with multiple comparisons (GraphPad Prism). A p-value of <0.05 was considered significant.

## Acknowledgements

Research reported in this publication was supported by the National Institute of Allergy and Infectious Diseases of the National Institutes of Health under awards R21AI166152 (to L.A.K. and J.R.E.) and F30AI169967 (to E.C.). J.R.E. is the Walter and Martha Renk Chair for Preharvest Food Safety. LAK holds an Investigators in the Pathogenesis of Infectious Disease Award from the Burroughs Wellcome Fund. The authors wish to thank the University of Wisconsin Translational Research Initiatives in Pathology (TRIP) Laboratory, supported by the University of Wisconsin Department of Pathology and Laboratory Medicine, UWCCC (P30 CA014520) and the Office of The Director of the National Institutes of Health (S10 OD023526), for its tissue processing and sectioning services. The funders had no role in study design, data collection and analysis, decision to publish, or preparation of the manuscript.

## Notes

### Competing Interest Statement

The authors have declared no competing interest.

## References

1. E. D. Weinberg, Nutritional Immunity: Host’s Attempt to Withhold Iron From Microbial Invaders. JAMA 231, 39–41 (1975).

2. C. C. Murdoch, E. P. Skaar, Nutritional immunity: the battle for nutrient metals at the host–pathogen interface. Nat Rev Microbiol 20, 657–670 (2022).

3. T. Goswami, et al., Natural-resistance-associated macrophage protein 1 is an H+/bivalent cation antiporter. Biochem J 354, 511–519 (2001).

4. H. Gunshin, et al., Cloning and characterization of a mammalian proton-coupled metal-ion transporter. Nature 388, 482–488 (1997).

5. B. Mackenzie, M. L. Ujwal, M.-H. Chang, M. F. Romero, M. A. Hediger, Divalent metal-ion transporter DMT1 mediates both H+ -coupled Fe2+ transport and uncoupled fluxes. Pflugers Arch 451, 544–558 (2006).

6. A. C. Illing, A. Shawki, C. L. Cunningham, B. Mackenzie, Substrate profile and metal-ion selectivity of human divalent metal-ion transporter-1. J Biol Chem 287, 30485–30496 (2012).

7. H. Fujishiro, Y. Yano, Y. Takada, M. Tanihara, S. Himeno, Roles of ZIP8, ZIP14, and DMT1 in transport of cadmium and manganese in mouse kidney proximal tubule cells. Metallomics 4, 700–708 (2012).

8. M. D. Fleming, et al., Nramp2 is mutated in the anemic Belgrade (b) rat: Evidence of a role for Nramp2 in endosomal iron transport. Proceedings of the National Academy of Sciences 95, 1148–1153 (1998).

9. J. Plant, A. A. Glynn, Natural resistance to Salmonella infection, delayed hypersensitivity and Ir genes in different strains of mice. Nature 248, 345–347 (1974).

10. D. J. Bradley, Letter: Genetic control of natural resistance to Leishmania donovani. Nature 250, 353–354 (1974).

11. P. Gros, E. Skamene, A. Forget, Genetic control of natural resistance to Mycobacterium bovis (BCG) in mice. The Journal of Immunology 127, 2417–2421 (1981).

12. S. Vidal, et al., The Ity/Lsh/Bcg locus: natural resistance to infection with intracellular parasites is abrogated by disruption of the Nramp1 gene. J Exp Med 182, 655–666 (1995).

13. D. Malo, et al., Haplotype mapping and sequence analysis of the mouse Nramp gene predict susceptibility to infection with intracellular parasites. Genomics 23, 51–61 (1994).

14. G. Govoni, et al., The Bcg/Ity/Lsh locus: genetic transfer of resistance to infections in C57BL/6J mice transgenic for the Nramp1 Gly169 allele. Infect Immun 64, 2923–2929 (1996).

15. J. K. White, A. Stewart, J.-F. Popoff, S. Wilson, J. M. Blackwell, Incomplete glycosylation and defective intracellular targeting of mutant solute carrier family 11 member 1 (Slc11a1). Biochemical Journal 382, 811–819 (2004).

16. J. Hu, et al., Resistance to Salmonellosis in the Chicken Is Linked to NRAMP1 and TNC. Genome Res. 7, 693–704 (1997).

17. R. Bellamy, et al., Variations in the NRAMP1 gene and susceptibility to tuberculosis in West Africans. N Engl J Med 338, 640–644 (1998).

18. S. J. Meisner, et al., Association of NRAMP1 polymorphism with leprosy type but not susceptibility to leprosy per se in west Africans. Am J Trop Med Hyg 65, 733–735 (2001).

19. G. G. Braliou, P. I. Kontou, H. Boleti, P. G. Bagos, Susceptibility to leishmaniasis is affected by host SLC11A1 gene polymorphisms: a systematic review and meta-analysis. Parasitol Res 118, 2329–2342 (2019).

20. M. Cellier, et al., Expression of the human NRAMP1 gene in professional primary phagocytes: studies in blood cells and in HL-60 promyelocytic leukemia. J Leukoc Biol 61, 96–105 (1997).

21. F. Canonne-Hergaux, et al., Expression and subcellular localization of NRAMP1 in human neutrophil granules. Blood 100, 268–275 (2002).

22. Y. Valdez, et al., Nramp1 expression by dendritic cells modulates inflammatory responses during *Salmonella* Typhimurium infection. Cellular Microbiology 10, 1646–1661 (2008).

23. T. A. Nagy, S. M. Moreland, H. Andrews-Polymenis, C. S. Detweiler, The Ferric Enterobactin Transporter Fep Is Required for Persistent Salmonella enterica Serovar Typhimurium Infection. Infect Immun 81, 4063–4070 (2013).

24. O. Cunrath, D. Bumann, Host resistance factor SLC11A1 restricts Salmonella growth through magnesium deprivation. Science 366, 995–999 (2019).

25. A.-B. Blanc-Potard, E. A. Groisman, How Pathogens Feel and Overcome Magnesium Limitation When in Host Tissues. Trends in Microbiology 29, 98–106 (2021).

26. M. Nairz, et al., Slc11a1 limits intracellular growth of Salmonella enterica sv. Typhimurium by promoting macrophage immune effector functions and impairing bacterial iron acquisition. Cell Microbiol 11, 1365–1381 (2009).

27. G. Fritsche, et al., Nramp1 functionality increases inducible nitric oxide synthase transcription via stimulation of IFN regulatory factor 1 expression. J Immunol 171, 1994–1998 (2003).

28. G. Fritsche, M. Nairz, S. J. Libby, F. C. Fang, G. Weiss, Slc11a1 (Nramp1) impairs growth of Salmonella enterica serovar typhimurium in macrophages via stimulation of lipocalin-2 expression. J Leukoc Biol 92, 353–359 (2012).

29. J. F. Hedges, E. Kimmel, D. T. Snyder, M. Jerome, M. A. Jutila, Solute carrier 11A1 is expressed by innate lymphocytes and augments their activation. J Immunol 190, 4263–4273 (2013).

30. K. L. Lokken-Toyli, et al., Vitamin A deficiency impairs neutrophil-mediated control of Salmonella via SLC11A1 in mice. Nat Microbiol 9, 727–736 (2024).

31. M. D. Fleming, et al., Microcytic anaemia mice have a mutation in Nramp2, a candidate iron transporter gene. Nat Genet 16, 383–386 (1997).

32. A. C. Chua, E. H. Morgan, Manganese metabolism is impaired in the Belgrade laboratory rat. J Comp Physiol B 167, 361–369 (1997).

33. M. D. Garrick, et al., DMT1: A mammalian transporter for multiple metals. Biometals 16, 41–54 (2003).

34. J. A. Edwards, J. E. Hoke, Defect of intestinal mucosal iron uptake in mice with hereditary microcytic anemia. Proc Soc Exp Biol Med 141, 81–84 (1972).

35. F. Canonne-Hergaux, A.-S. Zhang, P. Ponka, P. Gros, Characterization of the iron transporter DMT1 (NRAMP2/DCT1) in red blood cells of normal and anemic mk/mkmice. Blood 98, 3823–3830 (2001).

36. M. A. Su, C. C. Trenor, J. C. Fleming, M. D. Fleming, N. C. Andrews, The G185R mutation disrupts function of the iron transporter Nramp2. Blood 92, 2157–2163 (1998).

37. M. Knöpfel, L. Zhao, M. D. Garrick, Transport of divalent transition-metal ions is lost in small-intestinal tissue of b/b Belgrade rats. Biochemistry 44, 3454–3465 (2005).

38. H. Gunshin, et al., Slc11a2 is required for intestinal iron absorption and erythropoiesis but dispensable in placenta and liver. J Clin Invest 115, 1258–1266 (2005).

39. M. P. Mims, et al., Identification of a human mutation of DMT1 in a patient with microcytic anemia and iron overload. Blood 105, 1337–1342 (2005).

40. M. Priwitzerova, et al., Functional consequences of the human DMT1 (SLC11A2) mutation on protein expression and iron uptake. Blood 106, 3985–3987 (2005).

41. C. Beaumont, et al., Two new human DMT1 gene mutations in a patient with microcytic anemia, low ferritinemia, and liver iron overload. Blood 107, 4168–4170 (2006).

42. E. Bardou-Jacquet, et al., A novel N491S mutation in the human SLC11A2 gene impairs protein trafficking and in association with the G212V mutation leads to microcytic anemia and liver iron overload. Blood Cells Mol Dis 47, 243–248 (2011).

43. E. Blanco, C. Kannengiesser, B. Grandchamp, M. Tasso, C. Beaumont, Not all DMT1 mutations lead to iron overload. Blood Cells Mol Dis 43, 199–201 (2009).

44. W. Wu, et al., Divalent metal-ion transporter 1 is decreased in intestinal epithelial cells and contributes to the anemia in inflammatory bowel disease. Sci Rep 5, 16344 (2015).

45. J. Kim, et al., Influence of DMT1 and iron status on inflammatory responses in the lung. Am J Physiol Lung Cell Mol Physiol 300, L659–L665 (2011).

46. P. Urrutia, et al., Inflammation alters the expression of DMT1, FPN1 and hepcidin, and it causes iron accumulation in central nervous system cells. J Neurochem 126, 541–549 (2013).

47. R. M. Tsolis, M. N. Xavier, R. L. Santos, A. J. Bäumler, How to become a top model: impact of animal experimentation on human Salmonella disease research. Infect Immun 79, 1806–1814 (2011).

48. W. P. Loomis, et al., Temporal and Anatomical Host Resistance to Chronic Salmonella Infection Is Quantitatively Dictated by Nramp1 and Influenced by Host Genetic Background. PLoS One 9, e111763 (2014).

49. Y. Valdez, et al., Nramp1 drives an accelerated inflammatory response during Salmonella-induced colitis in mice. Cell Microbiol 11, 351–362 (2009).

50. L. F. Costa, T. A. Paixão, R. M. Tsolis, A. J. Bäumler, R. L. Santos, Salmonellosis in cattle: advantages of being an experimental model. Res Vet Sci 93, 1–6 (2012).

51. G. P. Ables, M. Nishibori, M. Kanemaki, T. Watanabe, Sequence analysis of the NRAMP1 genes from different bovine and buffalo breeds. J Vet Med Sci 64, 1081–1083 (2002).

52. J. Feng, et al., Bovine natural resistance associated macrophage protein 1 (Nramp1) gene. Genome Res. 6, 956–964 (1996).

53. W. J. Griffiths, A. L. Kelly, S. J. Smith, T. M. Cox, Localization of iron transport and regulatory proteins in human cells. QJM 93, 575–587 (2000).

54. H. Zoller, et al., Expression of the duodenal iron transporters divalent-metal transporter 1 and ferroportin 1 in iron deficiency and iron overload. Gastroenterology 120, 1412–1419 (2001).

55. Y. Harnik, et al., A spatial expression atlas of the adult human proximal small intestine. Nature 632, 1101–1109 (2024).

56. T. R. Powers, et al., Intracellular niche-specific profiling reveals transcriptional adaptations required for the cytosolic lifestyle of Salmonella enterica. PLOS Pathogens 17, e1009280 (2021).

57. K. Hantke, G. Nicholson, W. Rabsch, G. Winkelmann, Salmochelins, siderophores of Salmonella enterica and uropathogenic Escherichia coli strains, are recognized by the outer membrane receptor IroN. Proc Natl Acad Sci U S A 100, 3677–3682 (2003).

58. A. J. Bäumler, et al., IroN, a novel outer membrane siderophore receptor characteristic of Salmonella enterica. J Bacteriol 180, 1446–1453 (1998).

59. W. Rabsch, W. Voigt, R. Reissbrodt, R. M. Tsolis, A. J. Bäumler, Salmonella typhimurium IroN and FepA Proteins Mediate Uptake of Enterobactin but Differ in Their Specificity for Other Siderophores. J. Bacteriol. 181, 3610–3612 (1999).

60. D. G. Kehres, A. Janakiraman, J. M. Slauch, M. E. Maguire, SitABCD Is the Alkaline Mn2+ Transporter of Salmonella enterica Serovar Typhimurium. Journal of Bacteriology 184, 8 (2002).

61. V. E. Diaz-Ochoa, et al., *Salmonella* Mitigates Oxidative Stress and Thrives in the Inflamed Gut by Evading Calprotectin-Mediated Manganese Sequestration. Cell Host & Microbe 19, 814–825 (2016).

62. P. Petrarca, S. Ammendola, P. Pasquali, A. Battistoni, The Zur-Regulated ZinT Protein Is an Auxiliary Component of the High-Affinity ZnuABC Zinc Transporter That Facilitates Metal Recruitment during Severe Zinc Shortage. JB 192, 1553–1564 (2010).

63. R. L. Santos, S. Zhang, R. M. Tsolis, A. J. Bäumler, L. G. Adams, Morphologic and Molecular Characterization of Salmonella typhimurium Infection in Neonatal Calves. Vet Pathol 39, 200–215 (2002).

64. B. K. Coombes, et al., Analysis of the Contribution of Salmonella Pathogenicity Islands 1 and 2 to Enteric Disease Progression Using a Novel Bovine Ileal Loop Model and a Murine Model of Infectious Enterocolitis. Infect Immun 73, 7161–7169 (2005).

65. P. R. Watson, E. E. Galyov, S. M. Paulin, P. W. Jones, T. S. Wallis, Mutation of invH, but not stn, reduces Salmonella-induced enteritis in cattle. Infect Immun 66, 1432–1438 (1998).

66. J. R. Elfenbein, et al., Novel determinants of intestinal colonization of Salmonella enterica serotype typhimurium identified in bovine enteric infection. Infect Immun 81, 4311–4320 (2013).

67. S. Tandy, et al., Nramp2 expression is associated with pH-dependent iron uptake across the apical membrane of human intestinal Caco-2 cells. J Biol Chem 275, 1023–1029 (2000).

68. J. R. Forbes, P. Gros, Iron, manganese, and cobalt transport by Nramp1 (Slc11a1) and Nramp2 (Slc11a2) expressed at the plasma membrane. Blood 102, 1884–1892 (2003).

69. A. Shawki, et al., Intestinal DMT1 is critical for iron absorption in the mouse but is not required for the absorption of copper or manganese. Am J Physiol Gastrointest Liver Physiol 309, G635–647 (2015).

70. S. Gruenheid, M. Cellier, S. Vidal, P. Gros, Identification and characterization of a second mouse Nramp gene. Genomics 25, 514–525 (1995).

71. M. S. Madejczyk, N. Ballatori, The iron transporter ferroportin can also function as a manganese exporter. Biochimica et Biophysica Acta (BBA) - Biomembranes 1818, 651–657 (2012).

72. A. Donovan, et al., The iron exporter ferroportin/Slc40a1 is essential for iron homeostasis. Cell Metabolism 1, 191–200 (2005).

73. C. K. Fung, N. Zhao, The Combined Inactivation of Intestinal and Hepatic ZIP14 Exacerbates Manganese Overload in Mice. Int J Mol Sci 23, 6495 (2022).

74. K. Girijashanker, et al., Slc39a14 gene encodes ZIP14, a metal/bicarbonate symporter: similarities to the ZIP8 transporter. Mol Pharmacol 73, 1413–1423 (2008).

75. J. J. Pinilla-Tenas, et al., Zip14 is a complex broad-scope metal-ion transporter whose functional properties support roles in the cellular uptake of zinc and nontransferrin-bound iron. American Journal of Physiology-Cell Physiology 301, C862–C871 (2011).

76. J. P. Liuzzi, F. Aydemir, H. Nam, M. D. Knutson, R. J. Cousins, Zip14 (Slc39a14) mediates non-transferrin-bound iron uptake into cells. Proceedings of the National Academy of Sciences 103, 13612–13617 (2006).

77. I. S. Trowbridge, M. B. Omary, Human cell surface glycoprotein related to cell proliferation is the receptor for transferrin. Proc Natl Acad Sci U S A 78, 3039–3043 (1981).

78. M. Raffatellu, et al., Lipocalin-2 resistance confers an advantage to Salmonella enterica serotype Typhimurium for growth and survival in the inflamed intestine. Cell Host Microbe 5, 476–486 (2009).

79. S. Gruenheid, et al., The Iron Transport Protein NRAMP2 Is an Integral Membrane Glycoprotein That Colocalizes with Transferrin in Recycling Endosomes. Journal of Experimental Medicine 189, 831–841 (1999).

80. M. Tabuchi, T. Yoshimori, K. Yamaguchi, T. Yoshida, F. Kishi, Human NRAMP2/DMT1, Which Mediates Iron Transport across Endosomal Membranes, Is Localized to Late Endosomes and Lysosomes in HEp-2 Cells *. Journal of Biological Chemistry 275, 22220–22228 (2000).

81. N. Jabado, F. Canonne-Hergaux, S. Gruenheid, V. Picard, P. Gros, Iron transporter Nramp2/DMT-1 is associated with the membrane of phagosomes in macrophages and Sertoli cells. Blood 100, 2617–2622 (2002).

82. D. M. Johnson, S. Yamaji, J. Tennant, S. K. Srai, P. A. Sharp, Regulation of divalent metal transporter expression in human intestinal epithelial cells following exposure to non-haem iron. FEBS Letters 579, 1923–1929 (2005).

83. L. A. Knodler, et al., Dissemination of invasive Salmonella via bacterial-induced extrusion of mucosal epithelia. Proc. Natl. Acad. Sci. U.S.A. 107, 17733–17738 (2010).

84. L. A. Knodler, V. Nair, O. Steele-Mortimer, Quantitative assessment of cytosolic Salmonella in epithelial cells. PLoS ONE 9, e84681 (2014).

85. J. A. Klein, T. R. Powers, L. A. Knodler, Measurement of Salmonella enterica Internalization and Vacuole Lysis in Epithelial Cells. Methods Mol. Biol. 1519, 285–296 (2017).

86. M. L. Zaharik, B. A. Vallance, J. L. Puente, P. Gros, B. B. Finlay, Host-pathogen interactions: Host resistance factor Nramp1 up-regulates the expression of Salmonella pathogenicity island-2 virulence genes. Proc Natl Acad Sci U S A 99, 15705–15710 (2002).

87. N. Martin-Orozco, et al., Visualization of vacuolar acidification-induced transcription of genes of pathogens inside macrophages. Mol Biol Cell 17, 498–510 (2006).

88. A. Lane, et al., Sulfur- and phosphorus-standardized metal quantification of biological specimens using inductively coupled plasma mass spectrometry. STAR Protoc 3, 101334 (2022).

89. E. Gül, et al., *Salmonella* T3SS-2 virulence enhances gut-luminal colonization by enabling chemotaxis-dependent exploitation of intestinal inflammation. Cell Reports 43, 113925 (2024).

90. M. Barthel, et al., Pretreatment of mice with streptomycin provides a Salmonella enterica serovar Typhimurium colitis model that allows analysis of both pathogen and host. Infect Immun 71, 2839–2858 (2003).

91. J. Z. Liu, et al., Zinc Sequestration by the Neutrophil Protein Calprotectin Enhances *Salmonella* Growth in the Inflamed Gut. Cell Host & Microbe 11, 227–239 (2012).

92. D. E. Brown, et al., Increased Ferroportin-1 Expression and Rapid Splenic Iron Loss Occur with Anemia Caused by Salmonella enterica Serovar Typhimurium Infection in Mice. Infection and Immunity 83, 2290–2299 (2015).

93. K. Michels, E. Nemeth, T. Ganz, B. Mehrad, Hepcidin and Host Defense against Infectious Diseases. PLOS Pathogens 11, e1004998 (2015).

94. D.-K. Kim, et al., Inverse agonist of estrogen-related receptor γ controls Salmonella typhimurium infection by modulating host iron homeostasis. Nat Med 20, 419–424 (2014).

95. E. Deriu, et al., Probiotic Bacteria Reduce Salmonella Typhimurium Intestinal Colonization by Competing for Iron. Cell Host Microbe 14, 26–37 (2013).

96. M. Grander, et al., DMT1 Protects Macrophages from Salmonella Infection by Controlling Cellular Iron Turnover and Lipocalin 2 Expression. IJMS 23, 6789 (2022).

97. M. A. Dar, et al., Expression kinetics of natural resistance associated macrophage protein (NRAMP) genes in Salmonella Typhimurium-infected chicken. BMC Vet Res 14, 180 (2018).

98. M. A. Fischbach, et al., The pathogen-associated iroA gene cluster mediates bacterial evasion of lipocalin 2. Proc Natl Acad Sci U S A 103, 16502–16507 (2006).

99. R. R. Chaudhuri, et al., Comprehensive assignment of roles for Salmonella typhimurium genes in intestinal colonization of food-producing animals. PLoS Genet. 9, e1003456 (2013).

100. V. E. Diaz-Ochoa, et al., Salmonella Mitigates Oxidative Stress and Thrives in the Inflamed Gut by Evading Calprotectin-Mediated Manganese Sequestration. Cell Host & Microbe 19, 814–825 (2016).

101. J. Behnsen, et al., Siderophore-mediated zinc acquisition enhances enterobacterial colonization of the inflamed gut. Nat Commun 12, 7016 (2021).

102. L. F. Costa, et al., Iron acquisition pathways and colonization of the inflamed intestine by *Salmonella enterica* serovar Typhimurium. International Journal of Medical Microbiology 306, 604–610 (2016).

103. M. Cerasi, et al., The ZupT transporter plays an important role in zinc homeostasis and contributes to Salmonella enterica virulence. Metallomics 6, 845–853 (2014).

104. S. Ammendola, et al., High-Affinity Zn2+ Uptake System ZnuABC Is Required for Bacterial Zinc Homeostasis in Intracellular Environments and Contributes to the Virulence of Salmonella enterica. Infect Immun 75, 5867–5876 (2007).

105. M. Luckey, J. R. Pollack, R. Wayne, B. N. Ames, J. B. Neilands, Iron uptake in Salmonella typhimurium: utilization of exogenous siderochromes as iron carriers. J Bacteriol 111, 731–738 (1972).

106. C. N. Cornelissen, P. F. Sparling, Iron piracy: acquisition of transferrin-bound iron by bacterial pathogens. Mol Microbiol 14, 843–850 (1994).

107. M.-L. V. Crouch, M. Castor, J. E. Karlinsey, T. Kalhorn, F. C. Fang, Biosynthesis and IroC-dependent export of the siderophore salmochelin are essential for virulence of Salmonella enterica serovar Typhimurium. Molecular Microbiology 67, 971–983 (2008).

108. S. Yousuf, et al., Manganese import protects Salmonella enterica serovar Typhimurium against nitrosative stress. Metallomics 12, 1791–1801 (2020).

109. J. E. Karlinsey, M. E. Maguire, L. A. Becker, M. V. Crouch, F. C. Fang, The phage shock protein PspA facilitates divalent metal transport and is required for virulence of *Salmonella enterica* sv. Typhimurium. Molecular Microbiology 78, 669–685 (2010).

110. A. Janakiraman, J. M. Slauch, The putative iron transport system SitABCD encoded on SPI1 is required for full virulence of Salmonella typhimurium. Molecular Microbiology 35, 1146–1155 (2000).

111. W. J. Perry, et al., Staphylococcus aureus exhibits heterogeneous siderophore production within the vertebrate host. Proc Natl Acad Sci U S A 116, 21980–21982 (2019).

112. J. E. Cassat, et al., Integrated molecular imaging reveals tissue heterogeneity driving host-pathogen interactions. Sci Transl Med 10, eaan6361 (2018).

113. D. Galaris, A. Barbouti, K. Pantopoulos, Iron homeostasis and oxidative stress: An intimate relationship. Biochimica et Biophysica Acta (BBA) - Molecular Cell Research 1866, 118535 (2019).

114. J. D. Aguirre, V. C. Culotta, Battles with Iron: Manganese in Oxidative Stress Protection*. Journal of Biological Chemistry 287, 13541–13548 (2012).

115. L. He, K. Vasiliou, D. W. Nebert, Analysis and update of the human solute carrier (SLC) gene superfamily. Hum Genomics 3, 195 (2009).

116. M. A. Hediger, B. Clémençon, R. E. Burrier, E. A. Bruford, The ABCs of membrane transporters in health and disease (SLC series): Introduction. Molecular Aspects of Medicine 34, 95–107 (2013).

117. J. D. Rankin, R. J. Taylor, The estimation of doses of Salmonella typhimurium suitable for the experimental production of disease in calves. Vet Rec 78, 706–707 (1966).

118. P. W. Jones, P. Collins, M. M. Aitken, Passive protection of calves against experimental infection with Salmonella typhimurium. Vet Rec 123, 536–541 (1988).

119. L. A. Knodler, et al., Noncanonical inflammasome activation of caspase-4/caspase-11 mediates epithelial defenses against enteric bacterial pathogens. Cell Host Microbe 16, 249–256 (2014).

120. S. Porwollik, et al., Defined single-gene and multi-gene deletion mutant collections in Salmonella enterica sv Typhimurium. PLoS One 9, e99820 (2014).

121. P. P. Cherepanov, W. Wackernagel, Gene disruption in *Escherichia coli*: TcR and KmR cassettes with the option of Flp-catalyzed excision of the antibiotic-resistance determinant. Gene 158, 9–14 (1995).

122. J. A. Ibarra, et al., Induction of Salmonella pathogenicity island 1 under different growth conditions can affect Salmonella-host cell interactions in vitro. *Microbiology (Reading*, Engl.) 156, 1120–1133 (2010).

123. R. F. Wang, S. R. Kushner, Construction of versatile low-copy-number vectors for cloning, sequencing and gene expression in Escherichia coli. Gene 100, 195–199 (1991).

124. A. Holder, et al., Analysis of Genetic Variation in the Bovine SLC11A1 Gene, Its Influence on the Expression of NRAMP1 and Potential Association With Resistance to Bovine Tuberculosis. Front Microbiol 11, 1420 (2020).

125. A. P.-T. Jost, J. C. Waters, Designing a rigorous microscopy experiment: Validating methods and avoiding bias. J Cell Biol 218, 1452–1466 (2019).

126. R. C. Laughlin, et al., Spatial segregation of virulence gene expression during acute enteric infection with Salmonella enterica serovar Typhimurium. mBio 5, e00946–00913 (2014).

