## Supplemental Figures and Table for "SLC11A2 affects nutritional immunity in the gut epithelium"

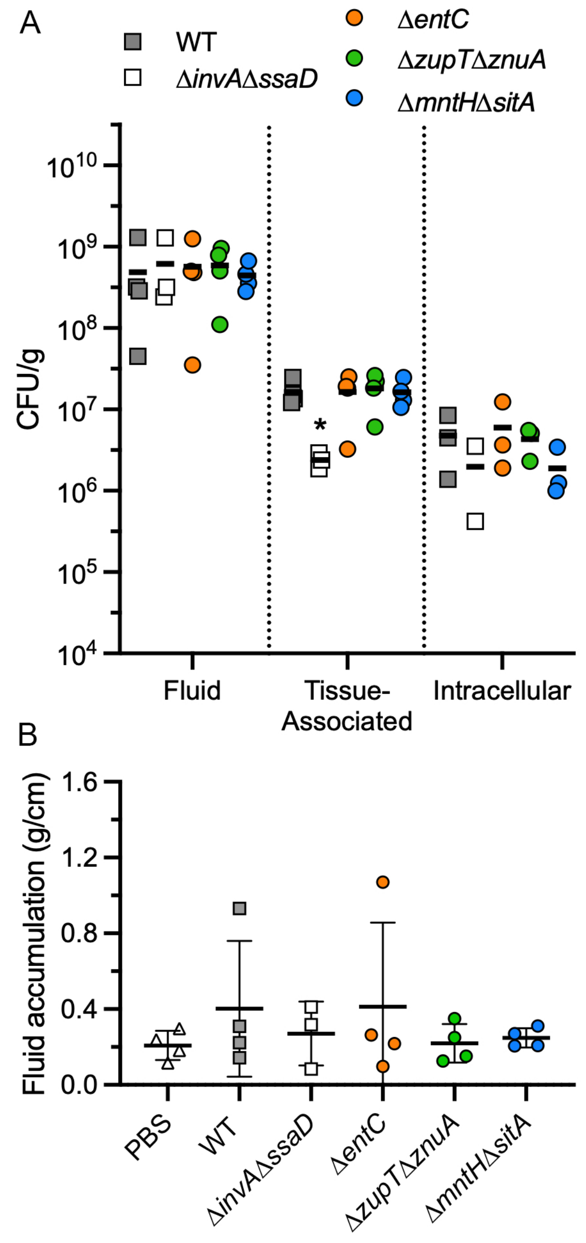


**Figure S1: Colonization and fluid accumulation at 2 h post-inoculation.** Ligated ileal loops were inoculated with PBS, wild-type (WT) bacteria, or the indicated deletion mutant (~$10^9$ CFU) for 2 h. (A) CFU from washed intestinal tissue (total tissue-asssicated bacteria), gentamicin-treated intestinal tissue (intracellular) and fluid were enumerated by serial dilution and plating on LB agar. CFU were noramlized to the tissue or fluid weight. (B) Fluid weight was nromalized to loop length. Each symbol represents data from one loop from one calf. *p<0.05, significqntly different from ST$/74 WT bacteria, ANOVA with multiple comparisons.

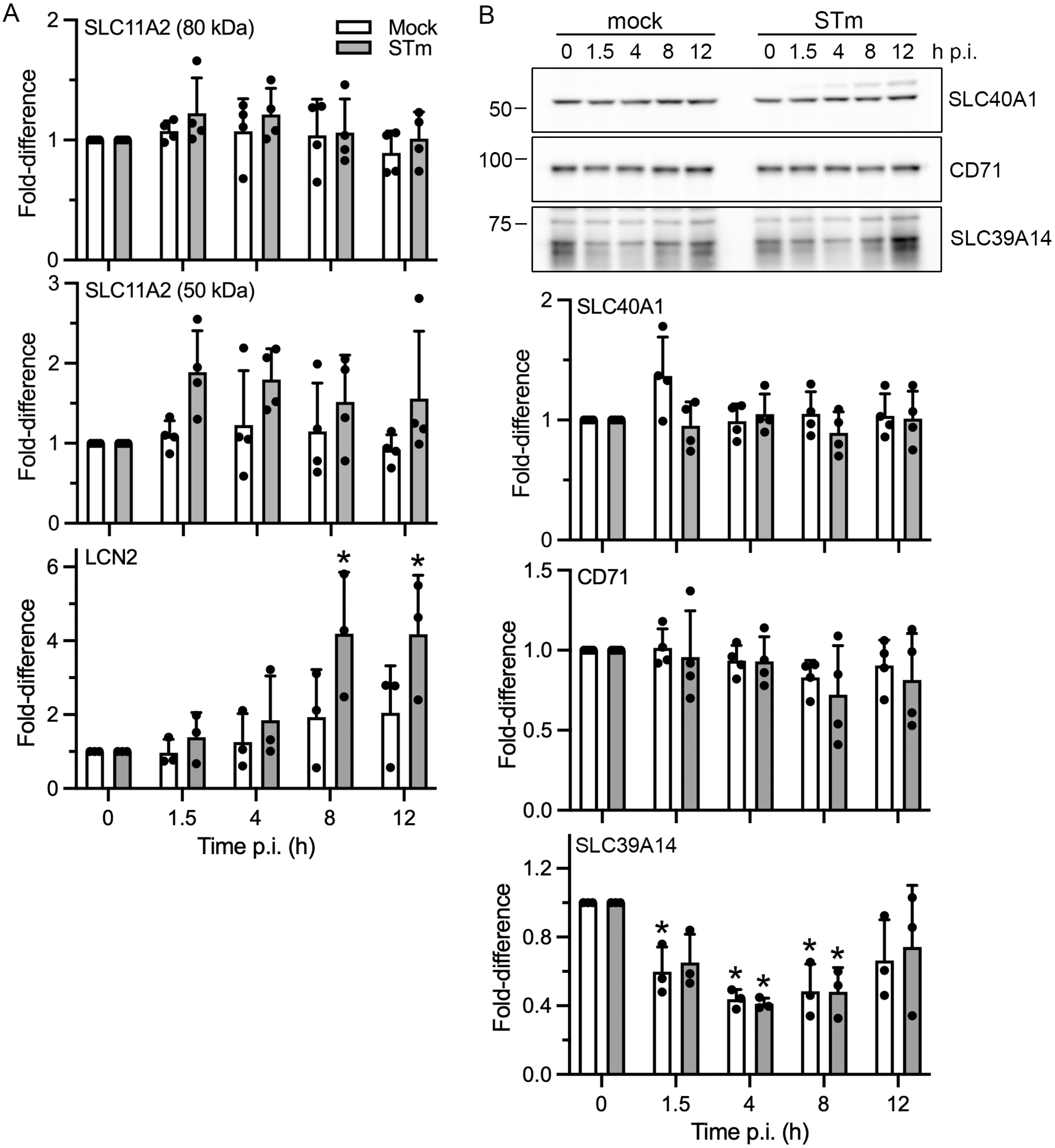


**Figure S2: Quantification of immunoblots from mock- and STm-infected HCT116 lysates.** Whole cell lysates were collected at the indicated times post-infection (p.i.) (A) Proteins were separated by SDS-PAGE and subject to immunoblotting with antibodies against SLC11A2 (NRAMP2), lipochalin-2 (LCN2), and cytochrome c oxidase IV (COX IV). Bands were quantified by densitometry (ImageJ) and normalized to COX IV (loading control) for each sample, then expressed as a fold-change compared to $t_0$. The 80 kDa and 50 kDa bands for SLC11A2, representing different isoforms, were quantified separately. (B) As for (A) except lysates were probed with antibodies against SLC40A1 (FPN1), CD71 (TfR) and SLC39A14 (ZIP14). Representative immunoblots are shown. Molecular mass markers are shown on the left. Bands were quantified by densitometry and normalized to COX IV (loading control) for each sample, then expressed as a fold-change compared to $t_0$. *$p<0.05$, ANOVA with Dunnett's post-hoc test. Mean ± SD from 3-4 independent experiments.

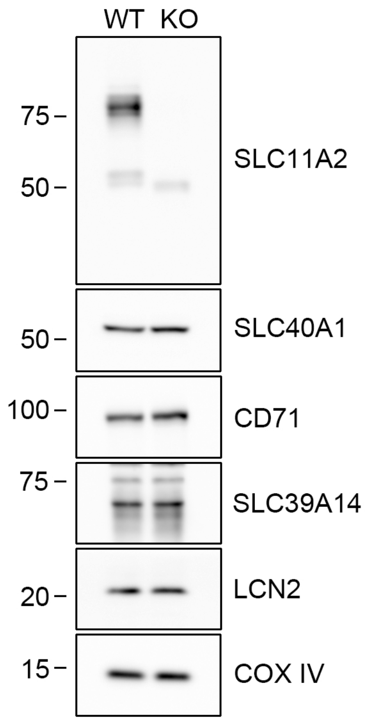


**Figure S3: Characterization of SLC11A2 knockout (KO) cells.** HCT116 SLC11A2 WT and KO cell lysates were probed with antibodies against SLC11A2, SLC40A1 (FPN1), CD71 (TfR), SLC39A14 (ZIP14), LCN2 and COXIV (loading control). Molecular mass markers are indicated on the left. Representative immunoblots are shown,

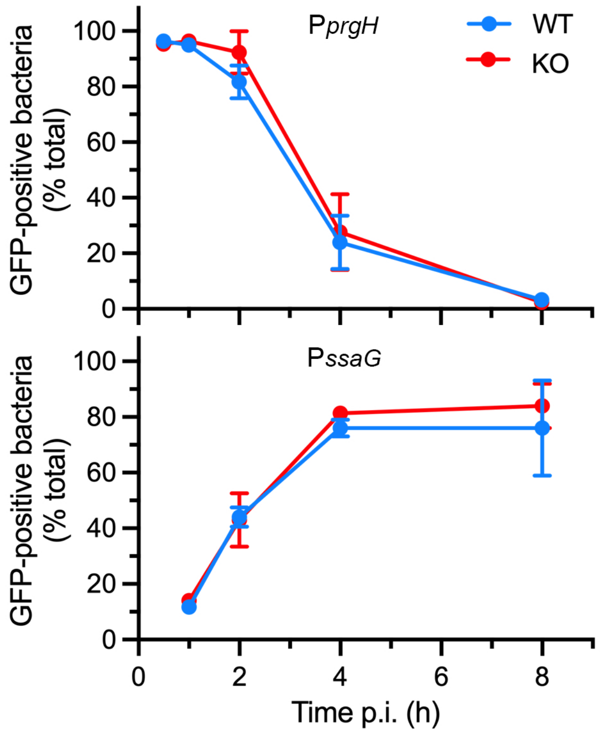


**Figure S4: Intracellular virulence gene expression.** HCT116 SLC11A2 WT and KO cells were infected with STM-mCherry harboring P*prgH-gfp(lva)* or P*ssaG-gfp(lva)* transcriptional reporters. At the times indicated, monolayers were fixed and the number of GFP-positive bacteria was scored by fluorescence microscopy. Mean ± SD from three independent experiments.

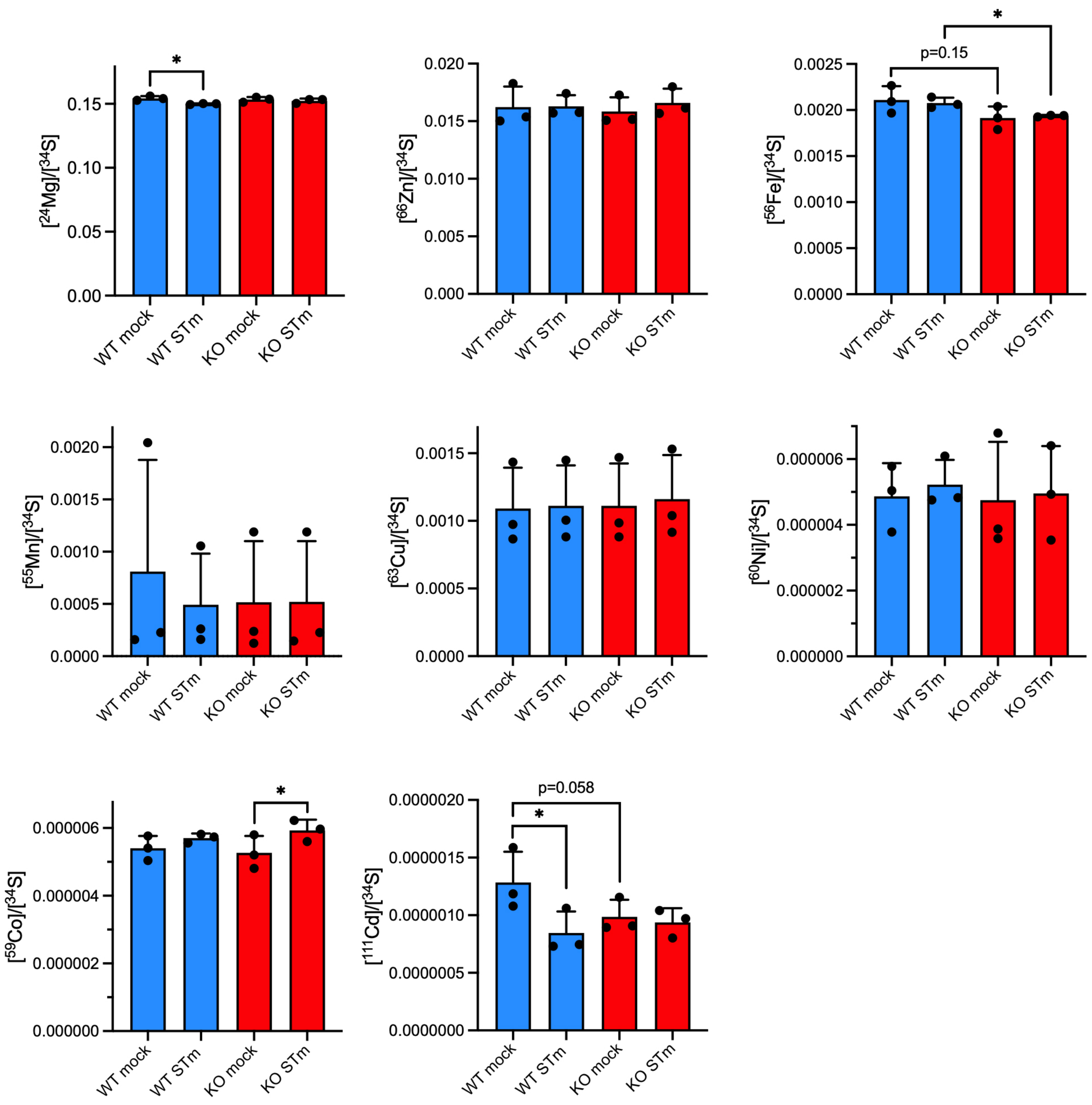


**Figure S5: ICP-MS characterization of SLC11A2 WT and KO cells.** HCT116 cells were mock- or STm-infected and collected at 8.h p.i. Total cellular levels of naturally occuring abundant isotopes of $^{24}Mg$, $^{66}Zn$, $^{56}Fe$, $^{55}Mn$, $^{63}Cu$, $^{60}Ni$, $^{59}Co$, $^{111}Cd$ and $^{34}S$ were quantified by ICP-MS analysis. The concentration of each metal (in ppb) was normalized to that of $^{34}S$ (in ppb) in each sample. Results are mean ± SD of three independent experiments. * $p<0.05$, Student's t-test.

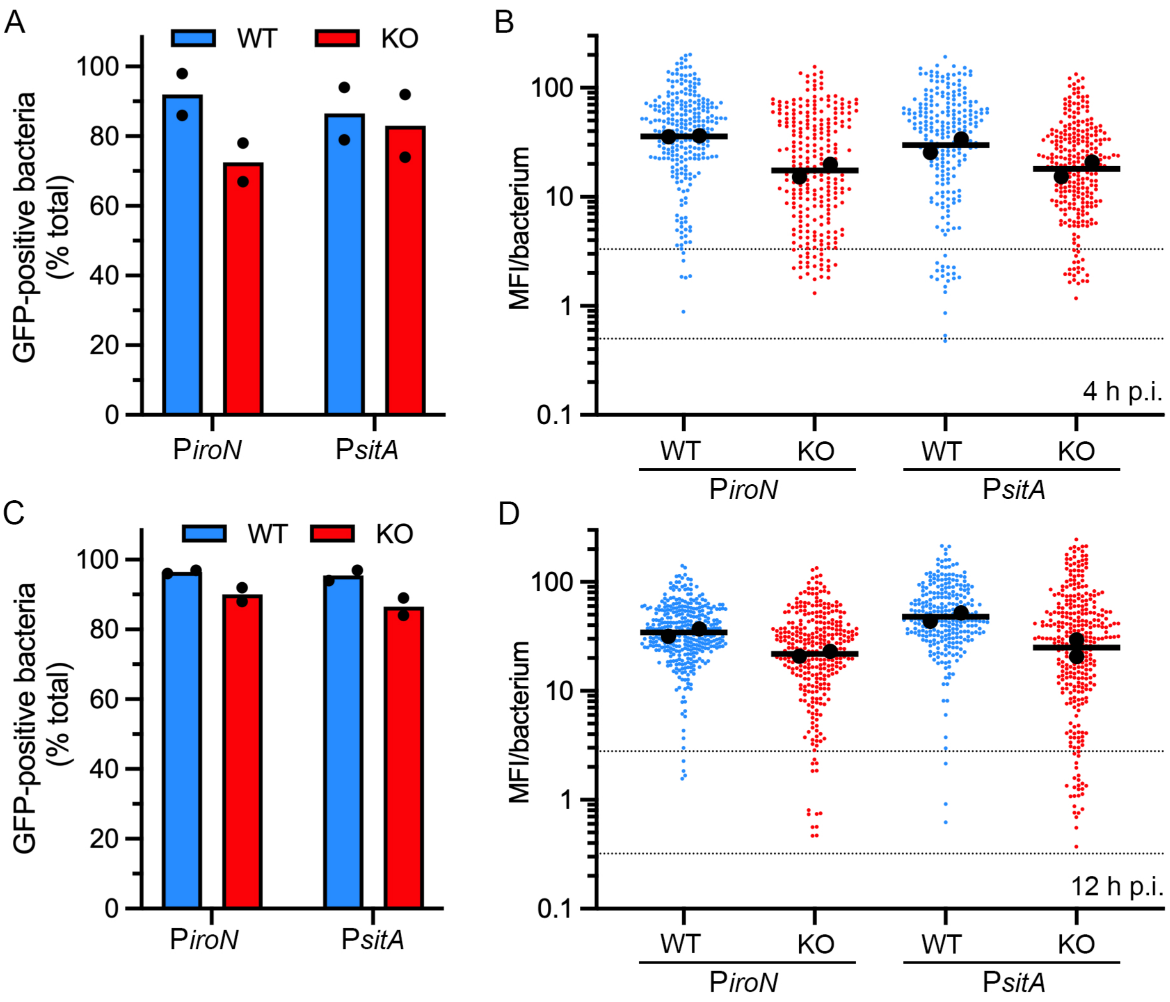


**Figure S6: SLC11A2 withholds iron and manganese from intracellular *Salmonella*.** (A and C) HCT116 SLC11A2 WT and KO cells seeded on glass coverslips were infected with STm-mCherry harboring P*iroN-gfpmut3* or P*sitA-gfpmut3* transcriptional reporters. At 4 h p.i. (A) and 12 h p.i. (C), monolayers were fixed and stained with Hoechst 33342 to label DNA. The number of GFP-positive bacteria was blindly scored by fluorescence microscopy. Mean from two independent experiments. (B and D) Quantification of the MFI signal at 4 h (B) and 12 h p.i.(D) by fluorescence microscopy and ImageJ analysis. Acquisition parameters (exposure time and gain) were the same for all reporters at each timepoint. Small dots represent individual bacteria; large dots indicate the mean of each experiment; horizontal bars indicate the average of two independent experiments. The dashed lines indicate the range of background fuorescence in the GFP channel for STm-mCherry (no reporter plasmid).

**Table S1: Oligonucleotides used for genetic complementation**

| **Name** | **Sequence (5' to 3")** | **Plasmid** |
|---|---|---|
| sitAcomp_XhoF | CCGCTCGAGGACCTGCCCAACGCATAATC | pWSK29-*sitA* |
| sitAcomp_HindR | CCCAAGCTTGAC**TCA**TTGTTGACTCCTCAG | pWSK29-*sitA* |
| mntHcomp_XhoF | CCGCTCGAGAACAAGTAACTGAATGACGT | pWSK29-*mntH* |
| mntHcomp_HindR | CCCAAGCT**TTA**TGACAACCCCATCACCG | pWSK29-*mntH* |

Engineered restriction sites are underlined. Stop codons are in **bold**.
